# Designing diverse and high-performance proteins with a large language model in the loop

**DOI:** 10.1101/2024.10.25.620340

**Authors:** Carlos A. Gomez-Uribe, Japheth Gado, Meiirbek Islamov

**Affiliations:** Solugen, Inc., Houston, TX, USA

## Abstract

We present a novel protein engineering approach to directed evolution with machine learning that integrates a new semi-supervised neural network fitness prediction model, Seq2Fitness, and an innovative optimization algorithm, **b**iphasic **a**nnealing for **d**iverse **a**daptive **s**equence **s**ampling (BADASS) to design sequences. Seq2Fitness leverages protein language models to predict fitness landscapes, combining evolutionary data with experimental labels, while BADASS efficiently explores these landscapes by dynamically adjusting temperature and mutation energies to prevent premature convergence and find diverse high-fitness sequences. Seq2Fitness predictions improve the Spearman correlation with fitness measurements over alternative model predictions, e.g., from 0.34 to 0.55 for sequences with mutations residues that are absent from the training set. BADASS requires less memory and computation compared to gradient-based Markov Chain Monte Carlo methods, while finding more higher-fitness sequences and maintaining sequence diversity in protein design tasks for two different protein families with hundreds of amino acids. For example, for both protein families 100% of the top 10,000 sequences found by BADASS have higher Seq2Fitness predictions than the wildtype sequence, versus a broad range between 3% to 99% for competing approaches with often many fewer than 10,000 sequences found. The fitness predictions for the top, top 100th, and top 1,000th sequences found by BADASS are all also higher. In addition, we developed a theoretical framework to explain where BADASS comes from, why it works, and how it behaves. Although we only evaluate BADASS here on amino acid sequences, it may be more broadly useful for exploration of other sequence spaces, including DNA and RNA. To ensure reproducibility and facilitate adoption, our code is publicly available [here](https://github.com/SoluLearn/BADASS/).

**Author summary:** Designing proteins with enhanced properties is essential for many applications, from industrial enzymes to therapeutic molecules. However, traditional protein engineering methods often fail to explore the vast sequence space effectively, partly due to the rarity of high-fitness sequences. In this work, we introduce BADASS, an optimization algorithm that samples sequences from a probability distribution with mutation energies and a temperature parameter that are updated dynamically, alternating between cooling and heating phases, to discover high-fitness proteins while maintaining sequence diversity. This stands in contrast to traditional approaches like simulated annealing, which often converge on fewer and lower fitness solutions, and gradient-based Markov Chain Monte Carlo (MCMC), also converging on lower fitness solutions and at a significantly higher computational and memory cost. Our approach requires only forward model evaluations and no gradient computations, enabling the rapid design of high-performing proteins that can be validated in the lab, especially when combined with our Seq2Fitness models. BADASS represents a significant advance in computational protein engineering, opening new possibilities for diverse applications.

## Introduction

Protein engineering plays a crucial role in biotechnology due to the transformative potential of high-performance proteins across a wide range of applications. Traditional approaches, such as directed evolution, are often time-consuming and labor-intensive, prone to becoming trapped in local optima, and limited to sequences with mostly a single mutation away from the starting sequence in each screening iteration [1]. In protein design, fitness is a quantitative description of the desired protein function, such as the conversion of substrate into product in a chemical reaction catalyzed by an enzyme. Recently, machine learning has been demonstrated to accelerate the discovery of proteins with improved fitness by overcoming limitations faced by traditional directed evolution through accurate prediction of fitness for sequences with multiple mutations, facilitating the in-silico exploration of broader regions of the sequence space [2–5].

Effective protein design with machine learning generally involves two key steps: first, building an accurate predictive model of protein fitness, and second, using this model to design a library of protein sequences that optimize the predicted fitness [6]. In recent years, protein language models have emerged as the state-of-the-art approach for predicting the effects of mutations on protein fitness [7,8]. However, zero-shot application of these models infers fitness from the distribution of amino acids in evolutionary data, which may diverge from experimentally measured or phenotypical fitness, particularly when the phenotypical fitness was not a target of evolutionary selection—a common occurrence in biotechnological applications [9]. Semi-supervised learning, integrating evolutionary predictions from zero-shot inference with experiment labels, has been shown to produce models that significantly improve the accuracy of phenotypical fitness prediction [10, 11].

Even with a fairly accurate model for predicting fitness, protein design efforts with machine learning may fail in the second step, if they do not produce high-fitness sequences that can be validated in the lab [6]. Generating a diverse set of high-fitness sequences in-silico maximizes the probability of finding proteins in the lab with the desired function and properties, such as stability or high expression levels. Since the sequence fitness landscape is discrete and vast, and predictive models are typically large and computationally expensive to evaluate, efficiently identifying high-scoring sequences, which are rare within the fitness landscape, is often a significant challenge [12]. Moreover, common optimization techniques, such as genetic algorithms and Markov chain Monte Carlo (MCMC) often struggle to efficiently navigate the vast sequence space [13, 14]. These challenges arise partly due to their intensive computational requirements, which limit the number of sequences explored under a fixed computational budget. As a result, the field is increasingly focusing on developing methods to identify diverse, high-fitness protein sequences using fitness models integrated within the optimization loop [15–17].

This paper presents a new approach for directed evolution with machine learning (Fig. 1). Our method integrates semi-supervised neural networks, named Seq2Fitness, which leverage protein language models to infer the fitness landscape from evolutionary density and experiment data. We also propose a novel protein sequence optimization algorithm–biphasic annealing for diverse adaptive sequence sampling (BADASS)–to design high-performance proteins with the Seq2Fitness or any other sequence-to-fitness machine learning models, requiring relatively few evaluations of the model without need for computing gradients. We compare our approach with the current alternatives described in [15] (EvoProtGrad) and [17] (GGS), demonstrating superior performance across design tasks using alpha-amylase (AMY BACSU) [18] and an endonuclease (NucB) [3]. We also developed a theory to motivate BADASS and explore why it works.

**Fig 1.**
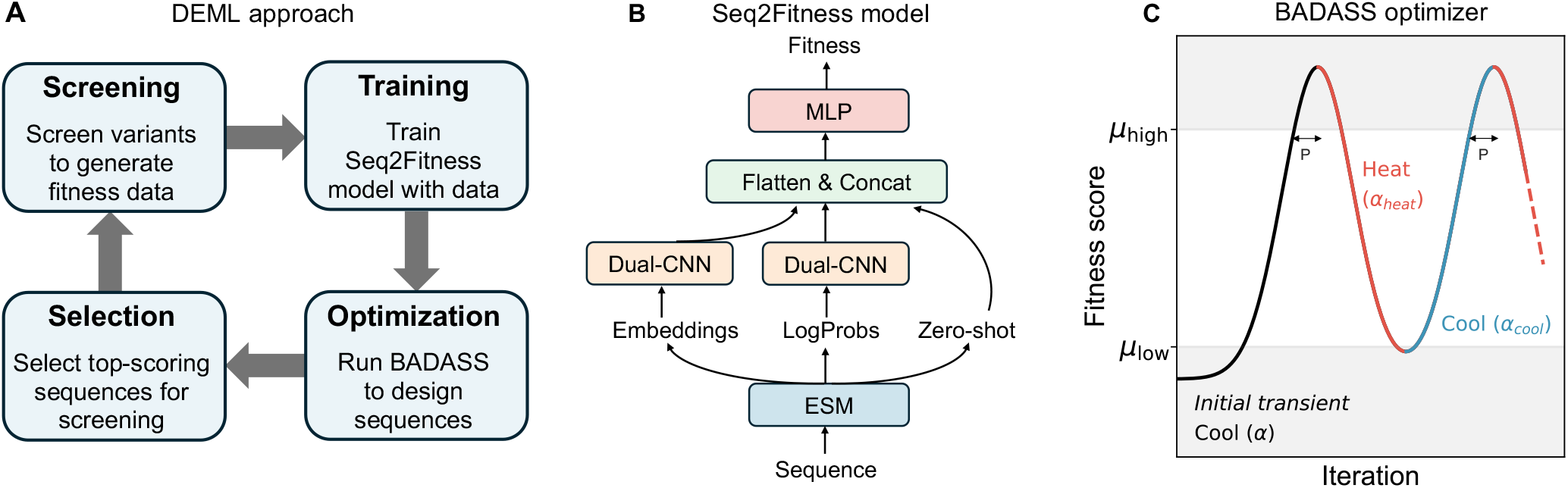
Overview of our approach. (A) Pipeline for directed evolution with machine learning using our proposed fitness prediction model and optimizer. New designs can start by running BADASS to select sequences with multiple mutations and high zero-shot scores from evolutionary models like ESM2 for an initial screening. (B) Architecture of the semi-supervised Seq2Fitness model. (C) The optimizer approach demonstrating the initial transient phase and two cooling and heating cycles. Sequences are iteratively sampled from an updated probability distribution, with the sampling temperature reduced (cooled) as the average fitness score rises. After the set point *µ*_high_ is breached, cooling continues for *patience* number of iterations, after which the temperature is increased (heated). The fitness score decreases until it reaches the set point *µ*_low_, at which point cooling resumes. The optimization process consists of multiple cooling and heating phases.

## Results

### Predicting protein fitness with Seq2Fitness

We developed a model, Seq2Fitness, to predict protein fitness from sequence. Seq2Fitness utilizes embeddings, log probabilities, and zero-shot scores from the ESM2-650M language model, and zero-shot scores from the ESM2-3B language model [19]. It employs parallel convolutional paths with novel statistical pooling layers to map sequence variants to experimental fitness measurements; see Fig. 1 and Materials and Methods section for more detail. We trained different Seq2Fitness models with published fitness datasets from the literature, including AAV [4], GB1 [20], NucB [3], and AMY BACSU [18], and compared the resulting models against alternative model architectures. We evaluated the models with different training/test dataset splits that collectively assess the ability of the model to extrapolate to new sequences, to new regions of the sequence space with a higher number of mutations, or to novel mutations beyond those seen in the training data. The dataset splits we used include (i) an 80/20 random sequence split, (ii) a two-vs-rest split [21], where all sequence variants with up to two mutations were included in the training set and the remainder in the testing set, and (iii) a mutational and (iv) a positional split [9, 22], where mutations or mutated positions in the test set were not present in any sequences in the training set.

Seq2Fitness consistently outperformed the other models across dataset splits with superior average scores across the AAV, GB1, NucB and AMY BACSU datasets (Table 1). The results on individual data splits showed that the improvement in performance of Seq2Fitness was most pronounced when extrapolating to new mutations and positions not present in the training set. Specifically, Seq2Fitness achieved average scores of 0.72 and 0.55 on mutational and positional splits, respectively, compared to 0.59 and 0.34 for the next best models, representing a 22% and 64% improvement in scores, respectively. The results on individual datasets and splits are found in Tables 4-7 in S1 Appendix. Additionally, we found that removing components of the Seq2Fitness architecture led to decline in performance, on average, highlighting the importance of each component to overall performance. These results, in Tables 8-11 in S1 Appendix, underscore the ability of Seq2Fitness to learn the fitness landscape effectively, enabling it to make accurate generalizations for designing new sequences.

**Table 1.**
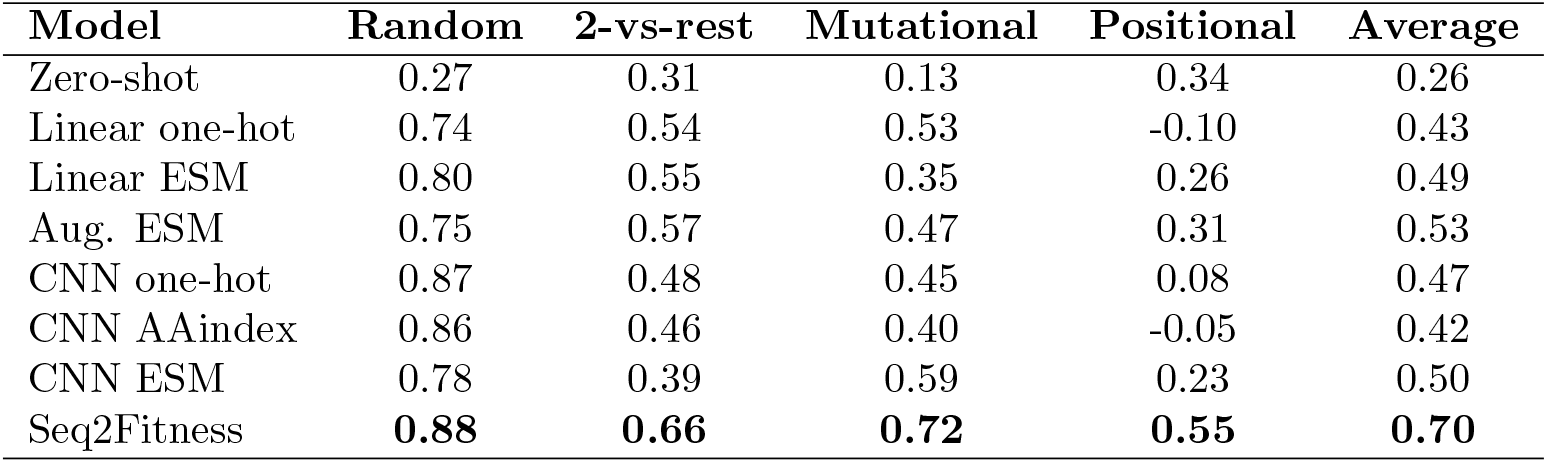
Performance comparison of Seq2Fitness and alternative models. Models were evaluated across different train/test splits, with performance metrics evaluated as Spearman correlation for regression tasks and adjusted AUC for classification tasks (NucB). Scores represent averages across the AAV, GB1, NucB, and AMY BACSU datasets.

### BADASS: Biphasic Annealing for Diverse Adaptive Sequence Sampling

We designed BADASS to efficiently explore the vast sequence landscape and identify diverse high-fitness variants. BADASS operates by iteratively sampling and scoring batches of sequences with a provided fitness model, such as Seq2Fitness. Each batch of sequences is sampled from a probability distribution that is dynamically updated based on mutation energies computed from the scores of previously evaluated batches, and a temperature parameter that is adjusted as the optimization progresses. In contrast to traditional simulated annealing [23, 24]–which gradually cools the system but leaves the energy function constant, often resulting in fewer high-scoring sequences and reduced diversity, due to a rapid decline in fitness score variance (Fig. 5)–BADASS utilizes dynamic temperature control to sustain a high score variance throughout the optimization process. As the optimization progresses, BADASS typically results in oscillations between regions with low and high scores across iterations. The mutation energies are also updated at every iteration, resulting in a dynamic approach that prevents premature convergence and promotes the discovery of more diverse sequences with higher scores. We developed theory to explain why the particular combination of dynamic temperature and mutation energy adjustments, along with the specific form of the mutation energies, makes BADASS an effective optimization approach to explore sequence space.

### Performance of BADASS in protein optimization

We evaluated BADASS on protein design tasks, specifically to identify higher-scoring alpha-amylase (AMY BACSU) [18] and endonuclease (NucB) [3] sequences, with both ESM2-650M language model or Seq2Fitness predictions as the fitness score. Fig. 2 shows the average sequence score and its variability when exploring sequence space for the alpha-amylase tasks with BADASS. For each task, BADASS was benchmarked against EvoProtGrad [15] and GGS [17]. EvoProtGrad did not originally incorporate a temperature parameter for the case of single model, in effect defaulting to a temperature of 1.0. To improve EvoProtGrad’s performance, we also tested it with a temperature of 0.1, equivalent to the temperature value used in GGS for the otherwise identical MCMC sampling distribution. We also modified the code to ensure consistency across sequence scores obtained during the EvoProtGrad process and during re-evaluation with the same scoring model.

**Fig 2.**
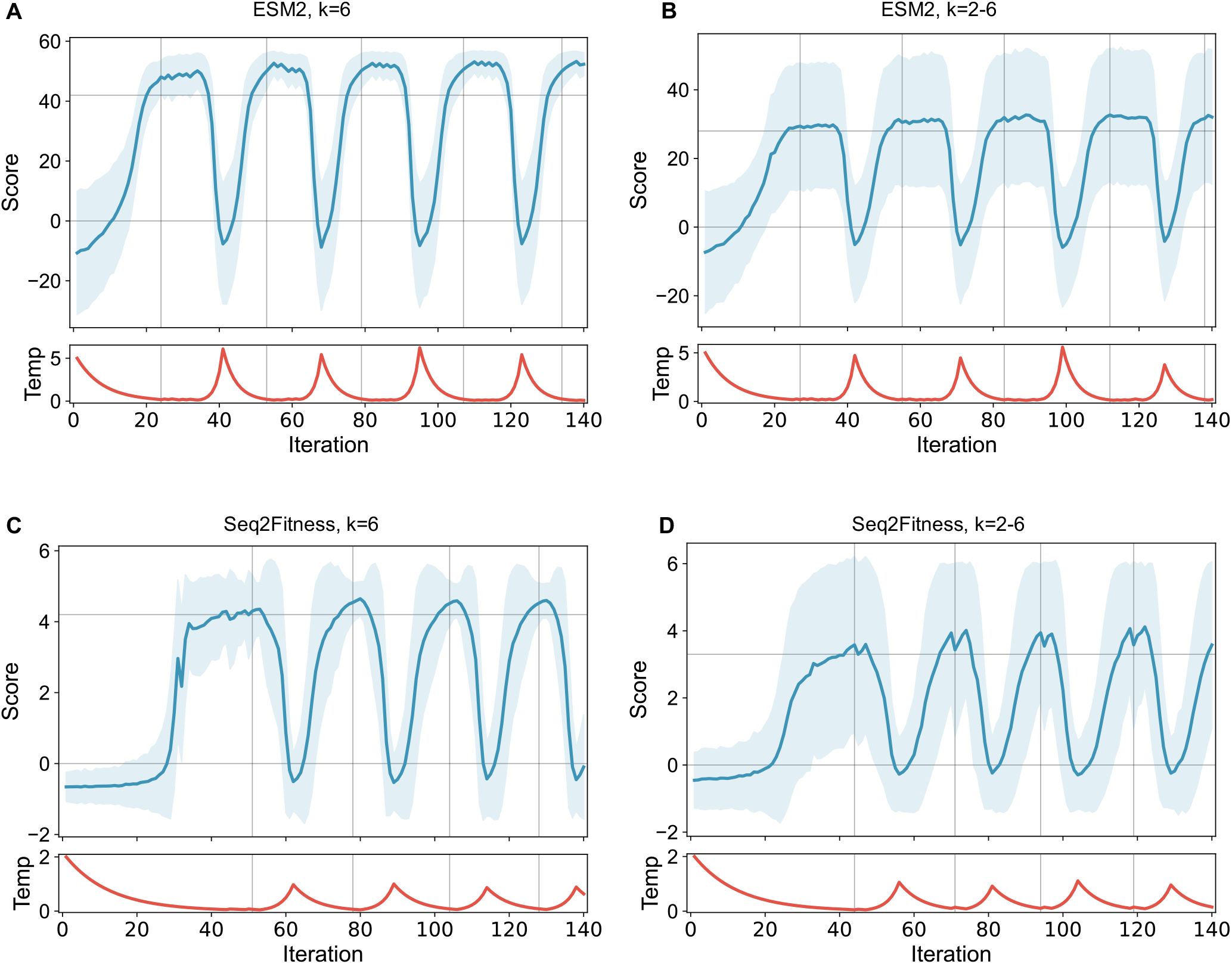
Fitness score statistics for BADASS optimization of alpha-amylase (AMY BACSU). BADASS was run for 140 iterations with a batch size of 1,000. The plot shows fitness score averaged over sampled sequences per iteration, with the shaded area representing scores within 1.96*σ* of the average, with *σ* the standard deviation of batch scores. Horizontal lines denote the set points *µ*_low_ and *µ*_high_ that govern the transitions between cooling and heating phases (Fig. 1C). Vertical dashed lines mark iterations where phase transitions occur as fitness scores cross the thresholds. The optimization was performed using the unsupervised ESM2 model and the semi-supervised Seq2Fitness model. Fitness scores were standardized as described in the methods. Runs included either exactly 6 mutations per variant (A, C) or an even mix of 2 to 6 mutations (B, D).

Table 2 summarizes the alpha-amylase optimization results, demonstrating that BADASS consistently outperformed EvoProtGrad in finding sequences with superior fitness scores. Specifically, 100% of the top 10,000 sequences generated by BADASS consistently achieved higher fitness scores than the reference sequence using both ESM2 and Seq2Fitness across sequence subspaces with different numbers of mutations *k*. In contrast, as few as 3.52% to 42.6% of the top 10,000 sequences found by EvoProtGrad are better than the reference sequence across temperatures and number of mutations for Seq2Fitness, and 90.1% to 99.5% for ESM2. Importantly, when using a temperature of 0.1 EvoProtGrad does not even find 10,000 unique sequences for this task or the NucB one. Moreover, applying GGS-smoothing with the Seq2Fitness model, as proposed by [17], did not clearly improve the results of either BADASS or EvoProtGrad sampling. E.g., GGS led to a higher proportion of sequences found being better than the wildtype sequence with EvoProtGrad, but a reduction in the same metric with BADASS, and the scores of the best, best 100th and best 1000th sequences relative to the corresponding Seq2Fitness EvoProtGrad or BADASS runs improved for some mutation numbers, remained unchanged in others, and decreased in others. Still, with GGS, BADASS outperformed EvoProtGrad with up to 73% of designed sequences having higher scores than the wildtype for BADASS, but only up to 44% for EvoProtGrad. Additionally, the best scoring sequences found by BADASS consistently achieved higher scores than the best sequences found by EvoProtGrad both with and without the GGS smoothing: BADASS found a sequence with an ESM2 score of 55.91 versus 54.04 for EvoProtGrad, and a sequence with a Seq2Fitness score of 5.97 versus 5.31 for EvoProtGrad.

**Table 2.** Alpha amylase sampling: Performance comparison between BADASS, EvoProtGrad and GGS (using EvoProtGrad on the Smoothed Seq2Fitness model) using ESM2 and Seq2Fitness models. All approaches are given comparable GPU compute time for the sampling; GGS requires an additional round to evaluate sequences with the original Seq2Fitness model. Metrics include the percentage of sequences better than wild type in the top 10,000 (or less when a method cannot find enough) sequences found, the best, best 100th, and best 1,000th sequence scores, and the number of unique mutations and unique mutated sites present in the top 10,000 sequences. The number of mutations per sequence is *k*. As benchmarks, the reference alpha amylase sequence has an ESM2 score of 0.0, and a Seq2Fitness score of 0.8. BADASS was run for 200 iterations with a batch size of 520 sequences. Missing entries for EvoProtGrad (using T=0.1 for ESM2) are due to the generation of a limited number of unique sequences (on the order of hundreds), as the sampler becomes overly concentrated on a small number of mutations.

| Fitness | Metric | BADASS |  |  | EvoProtGrad |  |  |  |
| --- | --- | --- | --- | --- | --- | --- | --- | --- |
|  |  | Gamma = 1 |  |  | T = 0.1 |  | T = 1.0 |  |
|  |  | k = 2-6 | 2 | 6 | 1-2 | 1-6 | 1-2 | 1-6 |
| **ESM2** | % better than WT | **100** | **100** | **100** | 97.67 | 99.52 | 90.10 | 97.00 |
|  | Best score | **55.91** | 20.19 | 54.12 | 18.80 | 54.04 | 18.57 | 45.60 |
|  | 100<sup>th</sup> best score | **52.24** | 16.52 | 52.09 | - | 38.11 | 13.58 | 28.43 |
|  | 1000<sup>th</sup> best score | 44.10 | 14.33 | **50.28** | - | - | 6.83 | 17.27 |
|  | Unique mutations | 1418 | 1018 | 115 | 18 | 25 | 1743 | **1752** |
|  | Unique sites | 330 | 244 | 73 | 13 | 18 | **346** | 345 |
| **Seq2Fitness** | % better than WT | **100** | **100** | **100** | 36.32 | 42.60 | 7.33 | 3.52 |
|  | Best score | 5.64 | 5.22 | **5.97** | 4.84 | 5.31 | 4.84 | 3.18 |
|  | 100<sup>th</sup> best score | 5.19 | 4.72 | **5.51** | 0.91 | 1.31 | 0.96 | 0.90 |
|  | 1000<sup>th</sup> best score | 4.88 | 4.03 | **5.20** | - | - | 0.57 | 0.34 |
|  | Unique mutations | 655 | 3608 | 393 | 509 | 713 | 3167 | **3629** |
|  | Unique sites | 228 | 410 | 181 | 209 | 238 | 423 | **424** |
| **Smoothed**<br>**Seq2Fitness**<br>**(w/GGS)** | % better than WT | 49.00 | **72.97** | 33.12 | 41.49 | 44.40 | 8.32 | 3.68 |
|  | Best score | 4.70 | **5.07** | 4.39 | 4.84 | 4.84 | 4.84 | 3.18 |
|  | 100<sup>th</sup> best score | 2.43 | **4.15** | 2.95 | 0.92 | 1.23 | 0.97 | 0.86 |
|  | 1000<sup>th</sup> best score | 1.35 | **1.52** | 1.43 | - | - | 0.52 | 0.31 |
|  | Unique mutations | 3402 | 2735 | **3955** | 330 | 421 | 2744 | 3480 |
|  | Unique sites | 398 | 373 | 414 | 138 | 187 | 423 | **424** |

On the NucB optimization tasks (Table 3), BADASS similarly showed superior performance, with 100% of the top 10,000 sequences generated achieving higher scores than the reference sequence using both ESM2 and Seq2Fitness models. However, with EvoProtGrad, as few as 12.1% of top 10,000 sequences had higher fitness scores compared with the reference sequence. However, in contrast to alpha-amylase tasks, smoothing with GGS did not negatively affect the fraction of sequences that outperformed the reference sequence for NucB, and GGS led to higher fitness scores achieved by of the best scoring sequence. GGS also improved the performance of EvoProtGrad on both the fraction of high-scoring sequences and the fitness scores of the best sequences. Hence, we conclude that the advantage of GGS-smoothing may be dependent on the task and the nature of the fitness landscape of the specific protein under study. On this task, the best ESM2 score BADASS found was 45.10 versus 40.46 with EvoProtGrad, and the best Seq2Fitness score found was 6.24 with BADASS versus 4.62 with EvoProtGrad.

**Table 3.**
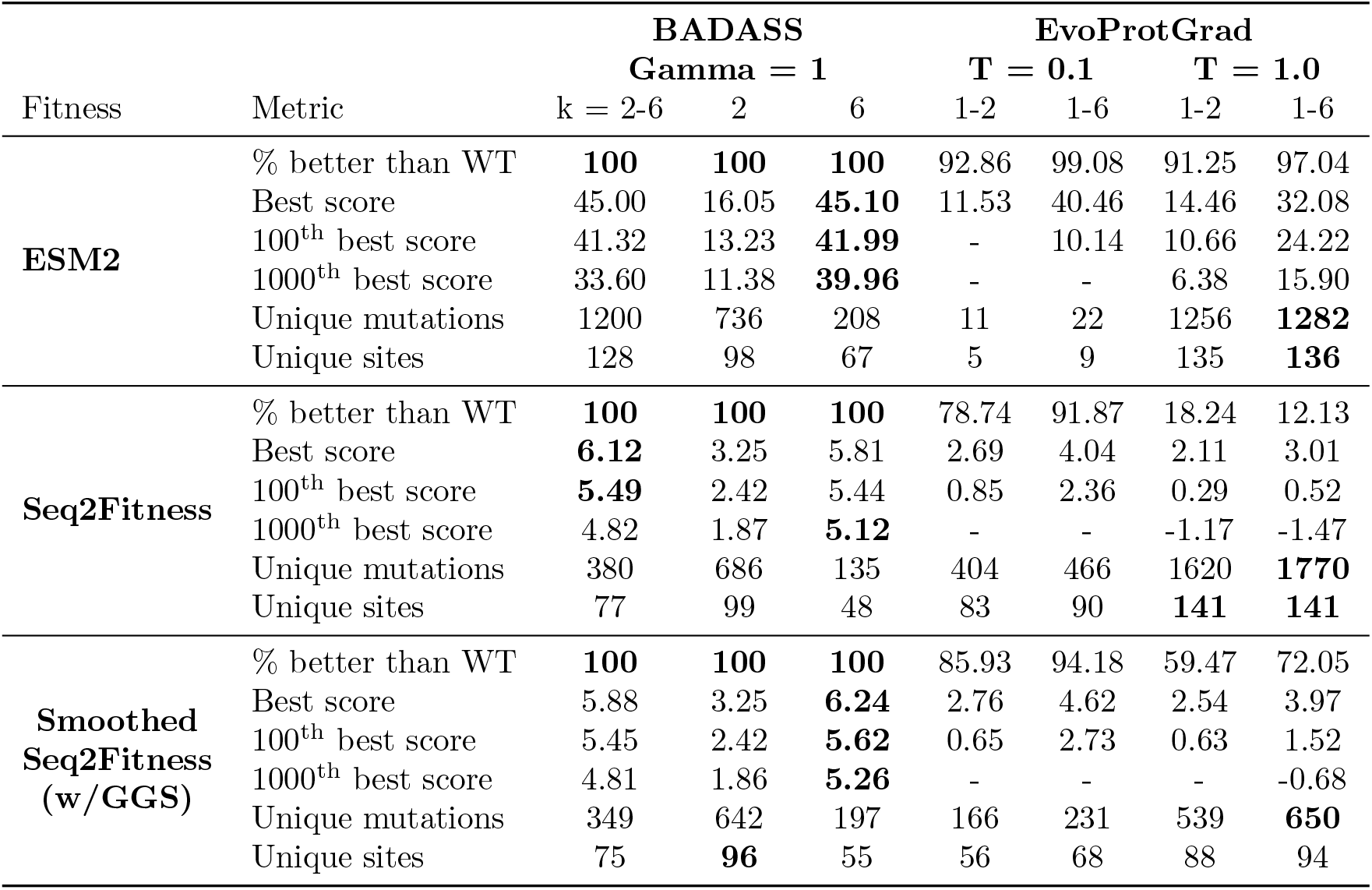
NucB sampling: Performance comparison between BADASS, EvoProtGrad and GGS using ESM2 and Seq2Fitness models. As benchmark, the reference NucB sequence has an ESM2 fitness of 0.0, and a Seq2Fitness score of -0.677. BADASS was run for 200 iterations with a batch size of 520 sequences.

**Table 4.** Performance of Seq2Fitness and alternative models with random split.

| Model | AAV | GB1 | AMY_BACSU | NucB | Average |
| --- | --- | --- | --- | --- | --- |
| Zero-shot | 0.05 | 0.08 | 0.61 | 0.33 | 0.27 |
| Linear one-hot | 0.80 | 0.79 | 0.65 | 0.71 | 0.74 |
| Linear ESM | 0.86 | 0.86 | 0.65 | 0.80 | 0.79 |
| Aug. linear | 0.81 | 0.79 | 0.65 | 0.77 | 0.75 |
| CNN one-hot | 0.91 | 0.92 | 0.72 | 0.92 | 0.87 |
| CNN AAindex | 0.91 | 0.92 | 0.72 | 0.92 | 0.87 |
| CNN ESM | **0.92** | 0.92 | 0.38 | 0.91 | 0.78 |
| Seq2Fitness | 0.91 | **0.93** | **0.75** | **0.92** | **0.88** |

**Table 5.** Performance of Seq2Fitness and alternative models with two-vs-rest split.

| Model | AAV | GB1 | AMY_BACSU | NucB | Average |
| --- | --- | --- | --- | --- | --- |
| Zero-shot | 0.15 | 0.11 | 0.54 | 0.45 | 0.31 |
| Linear one-hot | 0.66 | 0.57 | 0.57 | 0.37 | 0.54 |
| Linear ESM | 0.72 | 0.46 | 0.44 | 0.55 | 0.54 |
| Aug. linear | 0.66 | 0.58 | 0.58 | 0.46 | 0.57 |
| CNN one-hot | 0.69 | 0.53 | 0.23 | 0.49 | 0.48 |
| CNN AAindex | 0.51 | 0.32 | 0.07 | **0.66** | 0.39 |
| CNN ESM | 0.45 | 0.52 | 0.33 | 0.52 | 0.46 |
| Seq2Fitness | **0.77** | **0.62** | **0.61** | 0.62 | **0.66** |

**Table 6.** Performance of Seq2Fitness and alternative models with mutational split.

| Model | AAV | GB1 | AMY_BACSU | NucB | Average |
| --- | --- | --- | --- | --- | --- |
| Zero-shot | 0.19 | -0.39 | 0.62 | 0.10 | 0.13 |
| Linear one-hot | 0.61 | 0.57 | 0.52 | 0.42 | 0.53 |
| Linear ESM | 0.78 | 0.05 | 0.06 | 0.53 | 0.35 |
| Aug. linear | 0.61 | 0.21 | 0.62 | 0.46 | 0.47 |
| CNN one-hot | 0.74 | 0.48 | 0.34 | 0.23 | 0.45 |
| CNN AAindex | 0.84 | 0.58 | 0.28 | **0.68** | 0.59 |
| CNN ESM | 0.67 | 0.45 | 0.17 | 0.32 | 0.40 |
| Seq2Fitness | **0.82** | **0.69** | **0.71** | 0.64 | **0.72** |

**Table 7.** Performance of Seq2Fitness and alternative models with positional split. Positional splits with GB1 did not yield sufficient number of sequences for evaluation, and was, consequently, from the analyses.

| Model | AAV | GB1 | AMY_BACSU | NucB | Average |
| --- | --- | --- | --- | --- | --- |
| Zero-shot | 0.17 | NA | **0.66** | 0.20 | 0.34 |
| Linear one-hot | 0.01 | NA | 0.02 | -0.30 | -0.09 |
| Linear ESM | 0.26 | NA | 0.08 | 0.45 | 0.26 |
| Aug. linear | 0.11 | NA | 0.52 | 0.30 | 0.31 |
| CNN one-hot | 0.16 | NA | 0.35 | -0.28 | 0.08 |
| CNN AAindex | 0.45 | NA | 0.07 | 0.17 | 0.23 |
| CNN ESM | 0.10 | NA | 0.15 | -0.39 | -0.05 |
| Seq2Fitness | **0.53** | NA | 0.64 | **0.48** | **0.55** |

**Table 8.** Ablation of Seq2Fitness with random split.

| Model | AAV | GB1 | AMY_BACSU | NucB | Average |
| --- | --- | --- | --- | --- | --- |
| Seq2Fitness | **0.91** | **0.93** | **0.75** | **0.92** | **0.88** |
| w/ raw embeddings | 0.90 | 0.87 | 0.57 | 0.92 | 0.82 |
| w/o embeddings | 0.72 | 0.55 | 0.57 | 0.80 | 0.66 |
| w/o zero-shot scores | 0.91 | 0.92 | 0.73 | 0.92 | 0.87 |
| w/o normalized scores | 0.91 | 0.92 | 0.74 | 0.92 | 0.88 |
| w/o log-probabilities | 0.91 | 0.93 | 0.75 | 0.92 | 0.88 |

**Table 9.** Ablation of Seq2Fitness with two-vs-rest split.

| Model | AAV | GB1 | AMY_BACSU | NucB | Average |
| --- | --- | --- | --- | --- | --- |
| Seq2Fitness | 0.77 | **0.62** | **0.61** | 0.62 | **0.66** |
| w/ raw embeddings | 0.75 | 0.42 | 0.43 | 0.55 | 0.54 |
| w/o embeddings | 0.55 | 0.22 | 0.43 | 0.08 | 0.32 |
| w/o zero-shot scores | 0.73 | 0.61 | 0.58 | 0.64 | 0.64 |
| w/o normalized scores | 0.78 | 0.62 | 0.57 | **0.66** | 0.66 |
| w/o log-probabilities | **0.78** | 0.60 | 0.52 | 0.60 | 0.62 |

**Table 10.** Ablation of Seq2Fitness with mutational split.

| Model | AAV | GB1 | AMY_BACSU | NucB | Average |
| --- | --- | --- | --- | --- | --- |
| Seq2Fitness | **0.82** | **0.69** | **0.71** | 0.64 | **0.72** |
| w/ raw embeddings | 0.79 | 0.51 | 0.65 | **0.73** | 0.67 |
| w/o embeddings | 0.66 | -0.13 | 0.65 | 0.66 | 0.46 |
| w/o zero-shot scores | 0.80 | 0.67 | 0.70 | 0.62 | 0.70 |
| w/o normalized scores | 0.82 | 0.69 | 0.71 | 0.63 | 0.71 |
| w/o log-probabilities | 0.82 | 0.68 | 0.71 | 0.59 | 0.70 |

**Table 11.**
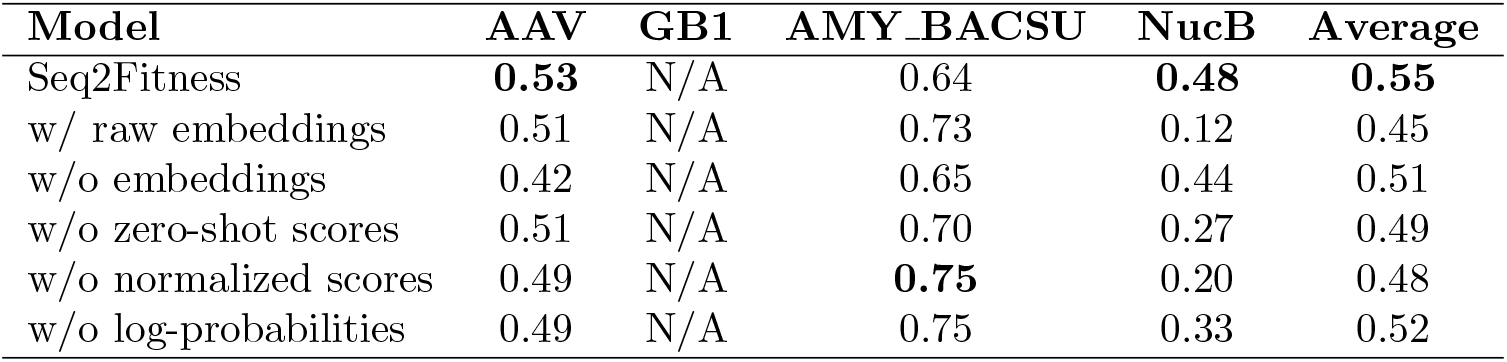
Ablation of Seq2Fitness with positional split.

| Model | AAV | GB1 | AMY_BACSU | NucB | Average |
| --- | --- | --- | --- | --- | --- |
| Seq2Fitness | **0.53** | N/A | 0.64 | **0.48** | **0.55** |
| w/ raw embeddings | 0.51 | N/A | 0.73 | 0.12 | 0.45 |
| w/o embeddings | 0.42 | N/A | 0.65 | 0.44 | 0.51 |
| w/o zero-shot scores | 0.51 | N/A | 0.70 | 0.27 | 0.49 |
| w/o normalized scores | 0.49 | N/A | **0.75** | 0.20 | 0.48 |
| w/o log-probabilities | 0.49 | N/A | 0.75 | 0.33 | 0.52 |

In addition to generating high-scoring sequences, BADASS simultaneously maintained a high level of diversity in the generated sequences despite achieving high fitness values. Paired with the Seq2Fitness model, BADASS identified sequences with a substantial number of unique mutations and mutated sites, representing up to 45% and 25% of possible mutations and 96% and 70% of possible sites in alpha-amylase and NucB respectively. In contrast, EvoProtGrad achieved similar levels of diversity with a high sampling temperature but at the expense of substantially lower fitness scores. EvoProtGrad, as with other MCMC approaches, tends to find sequences with high fitness scores only with a sufficiently low sampling temperature such that the diversity of generated sequences is low, as is evidenced by the number of unique mutations and unique sites. BADASS, however, is able to explore a broad range of sampling temperatures in a single run due to the dynamic temperature control and identify high scoring sequences without loss of diversity. Tables 12-15 in S2 Appendix show detailed results comparing BADASS and EvoProtGrad for alpha-amylase and NucB.

**Table 12.**
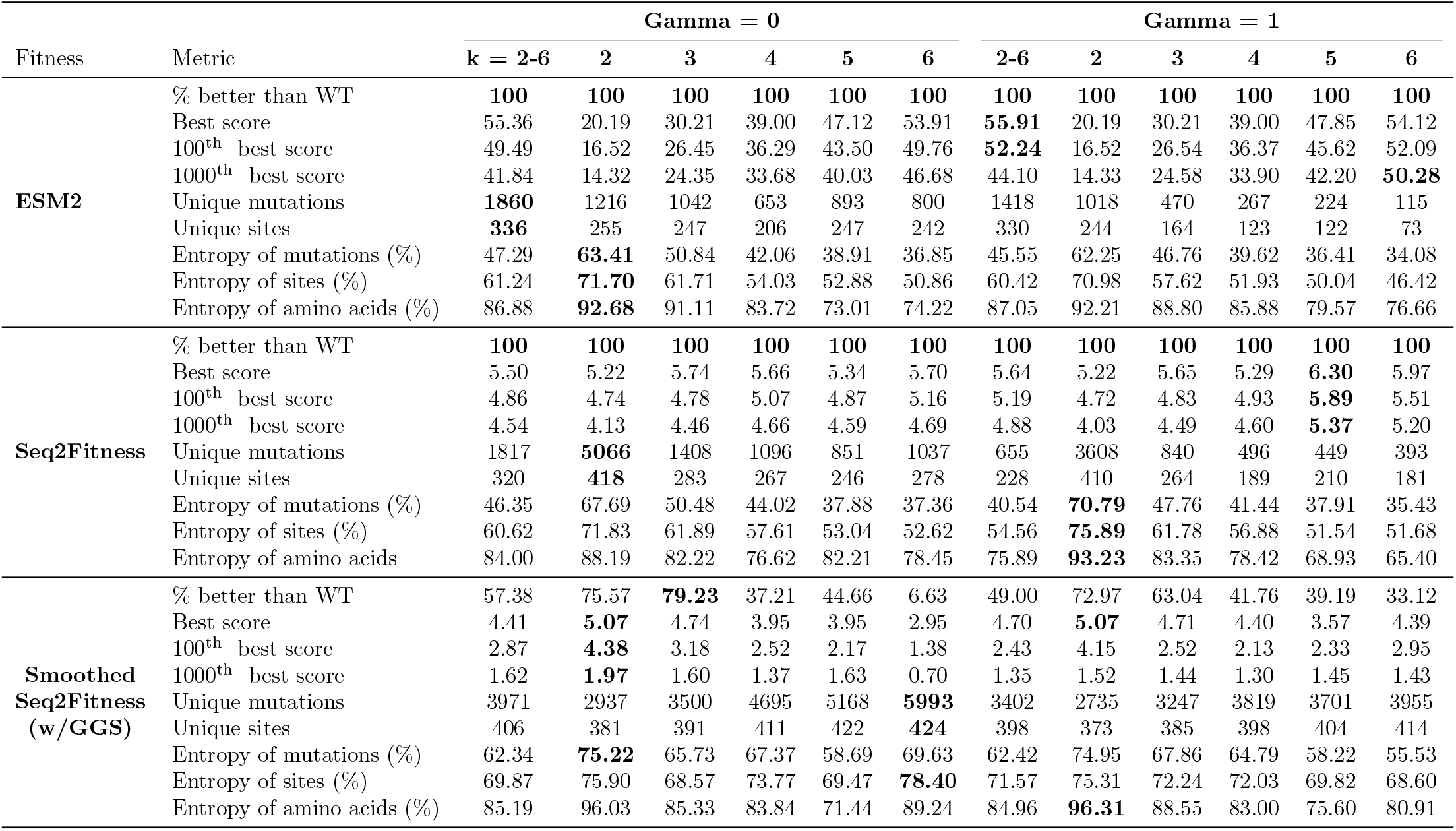
Fuller BADASS results on alpha amylase tasks. The entropies are shown as the percentage relative to the maximum entropy of a uniform distribution.

**Table 13.**
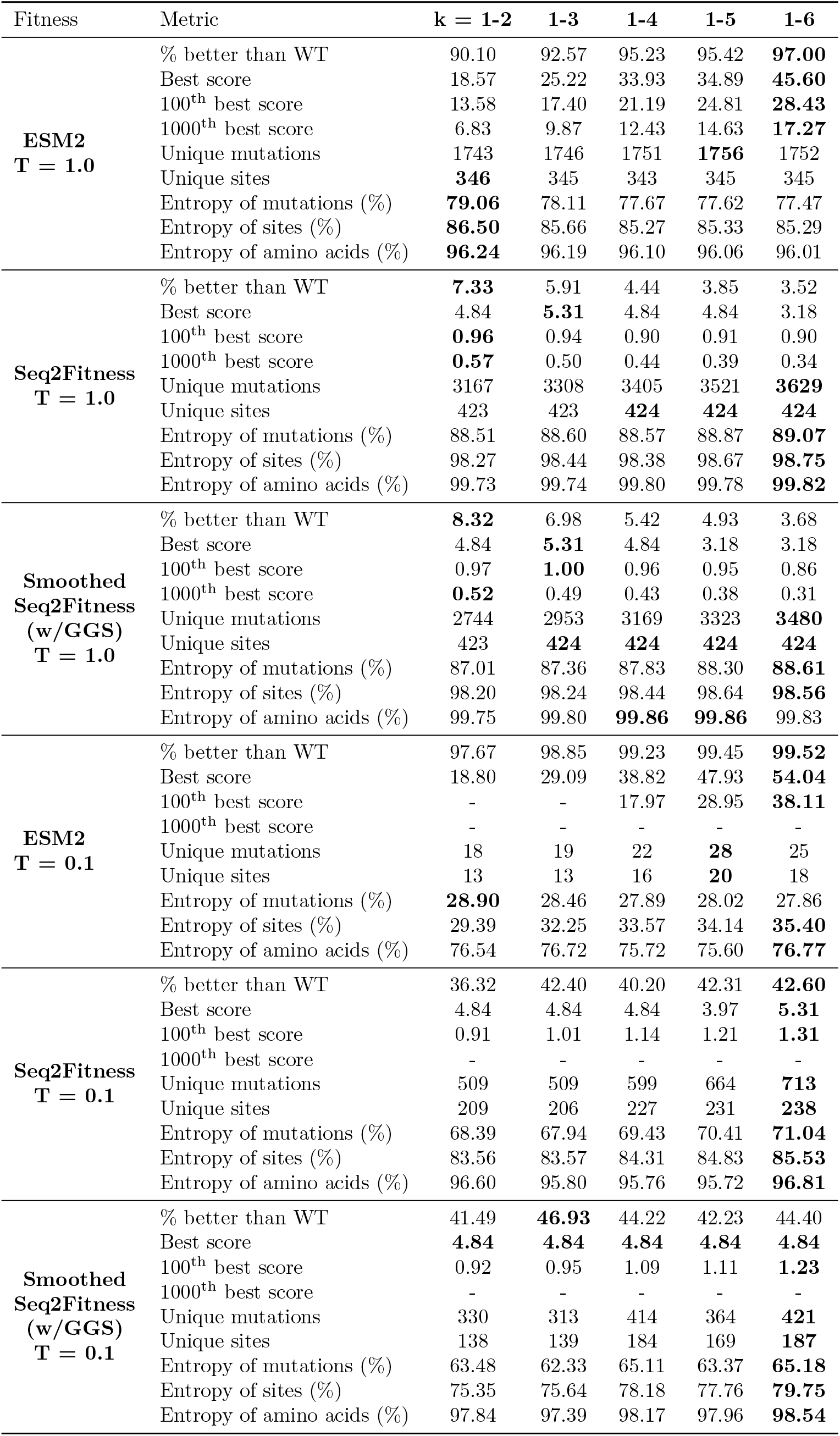
Fuller EvoProtGrad results on alpha amylase tasks (modified code).

Furthermore, we evaluated the contribution of the key features of BADASS, showing in Fig. 5 and Table 16 that replacing the temperature control based on the average score with a simple cooling schedule, or not updating the mutation energies based on the sequences sampled throughout the optimization, significantly hurts performance, designing only sequences with lower fitness scores. Taken together, these results demonstrate the effectiveness and robustness of the BADASS optimization algorithm for protein design.

**Table 14.**
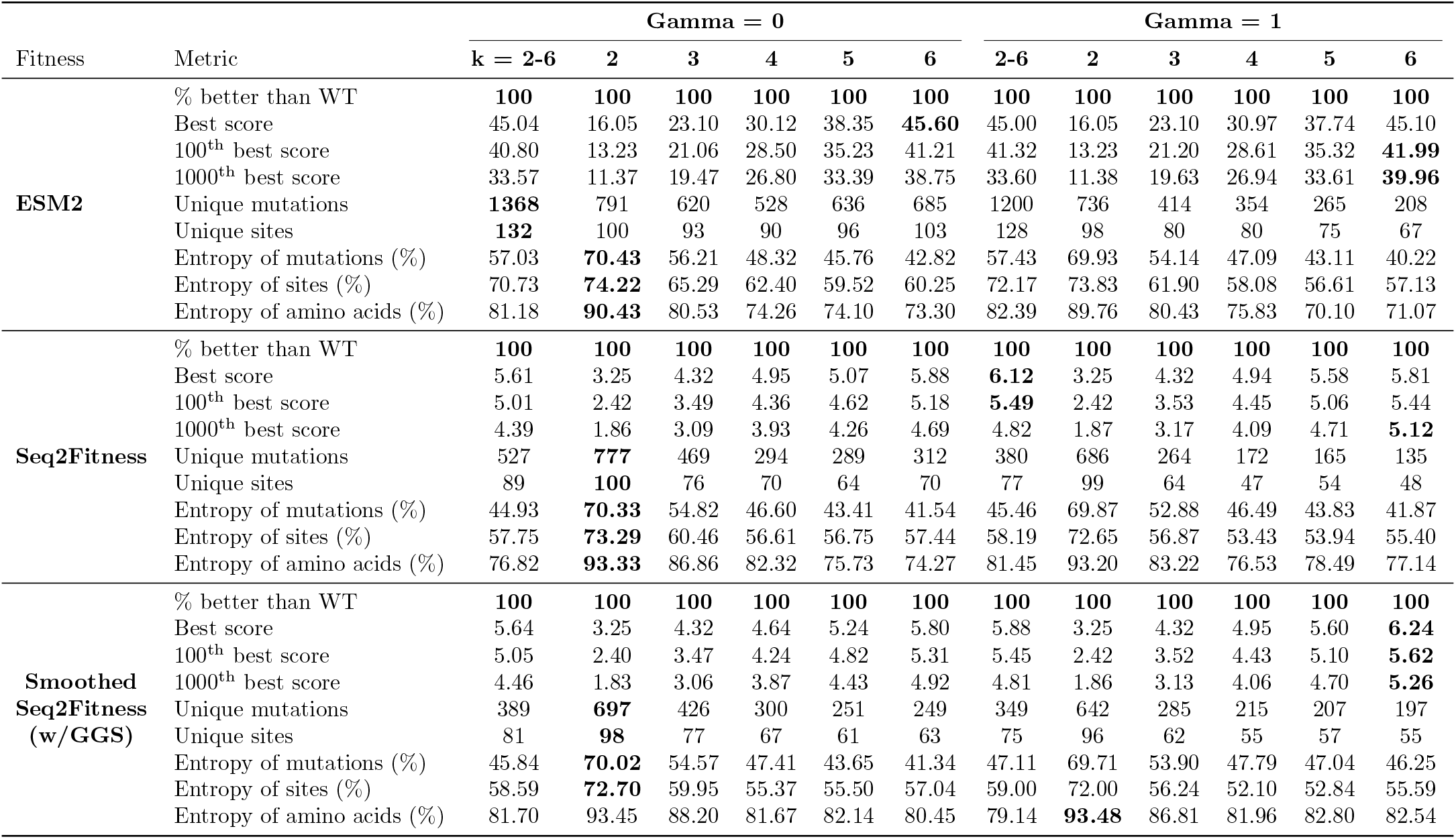
Fuller BADASS results on NucB tasks.

**Table 15.**
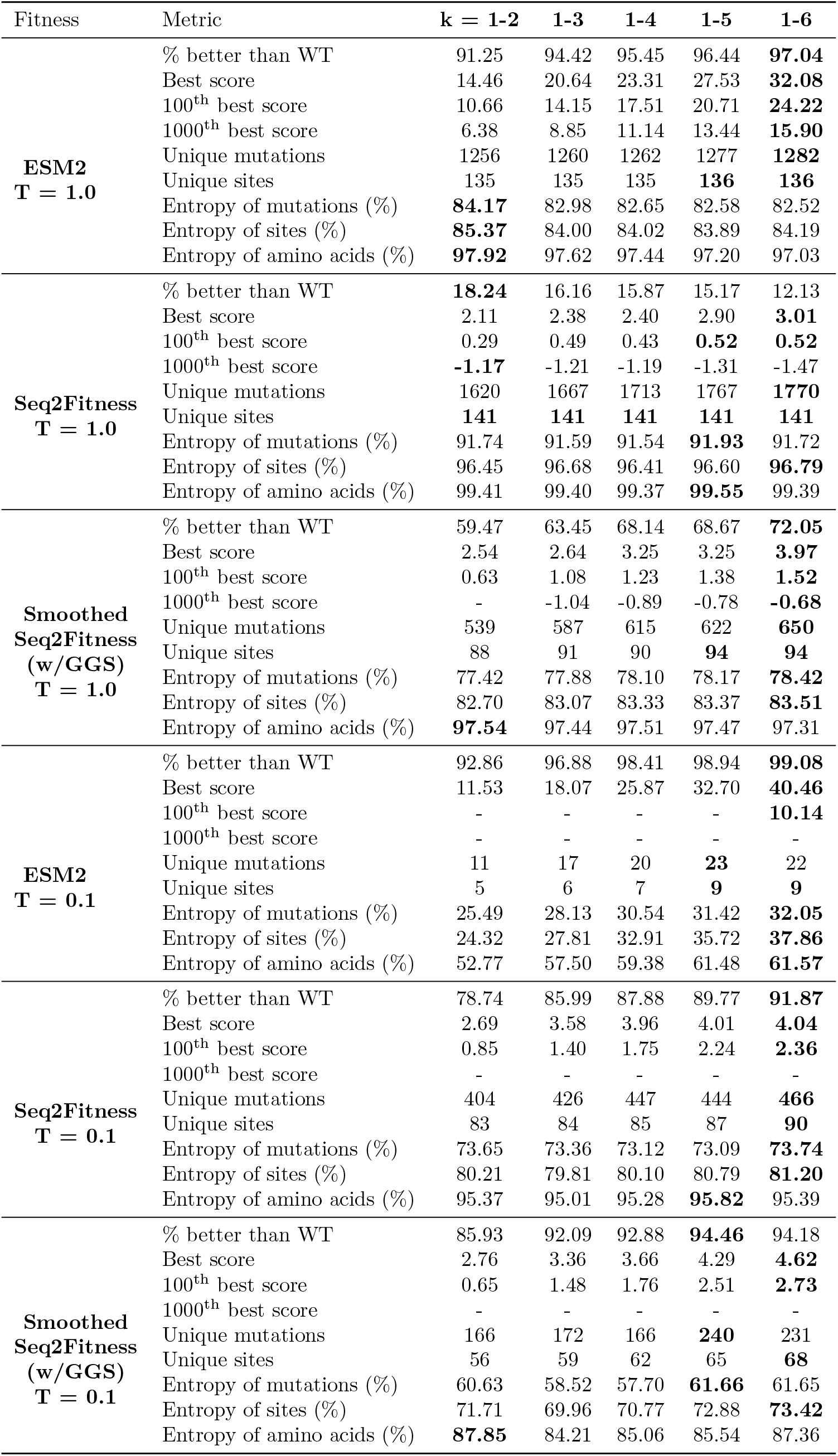
Fuller EvoProtGrad results on NucB tasks.

**Table 16.**
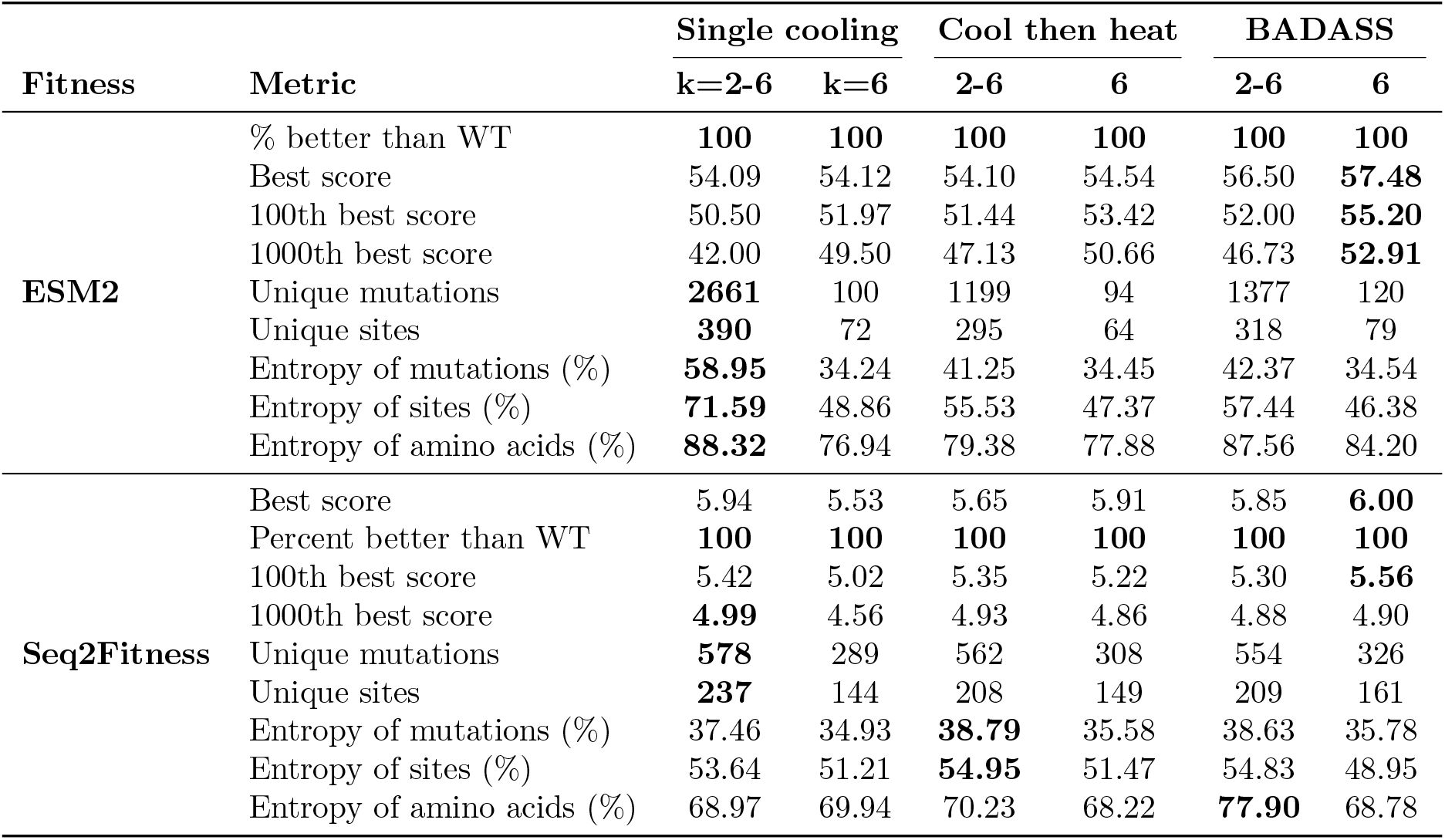
Comparison of BADASS with a simpler cooling and heating regime on alpha-amylase tasks, while still updating mutation energies in the sampler. The simpler cool then heat approach cools until the sampler is unable to find enough sequences for the batch, heats until a use-specified high temperature is reached, and then repeats. The BADASS controls find the highest scoring sequence for ESM2 and Seq2Fitness on the amylase task: 57.48 versus 54.54, and 6.00 versus 5.91, respectively. But the simpler temperature control is competitive, and requires a single user-specified set point (the high temperature) rather than two, so may be preferred in some cases. We also compare BADASS to having a single cooling schedule: BADASS finds better sequences. When the mutation energies are not updated, the resulting optimization gets stuck on sequences with mediocre scores, as seen clearly in Fig. 5.

**Table 17.**
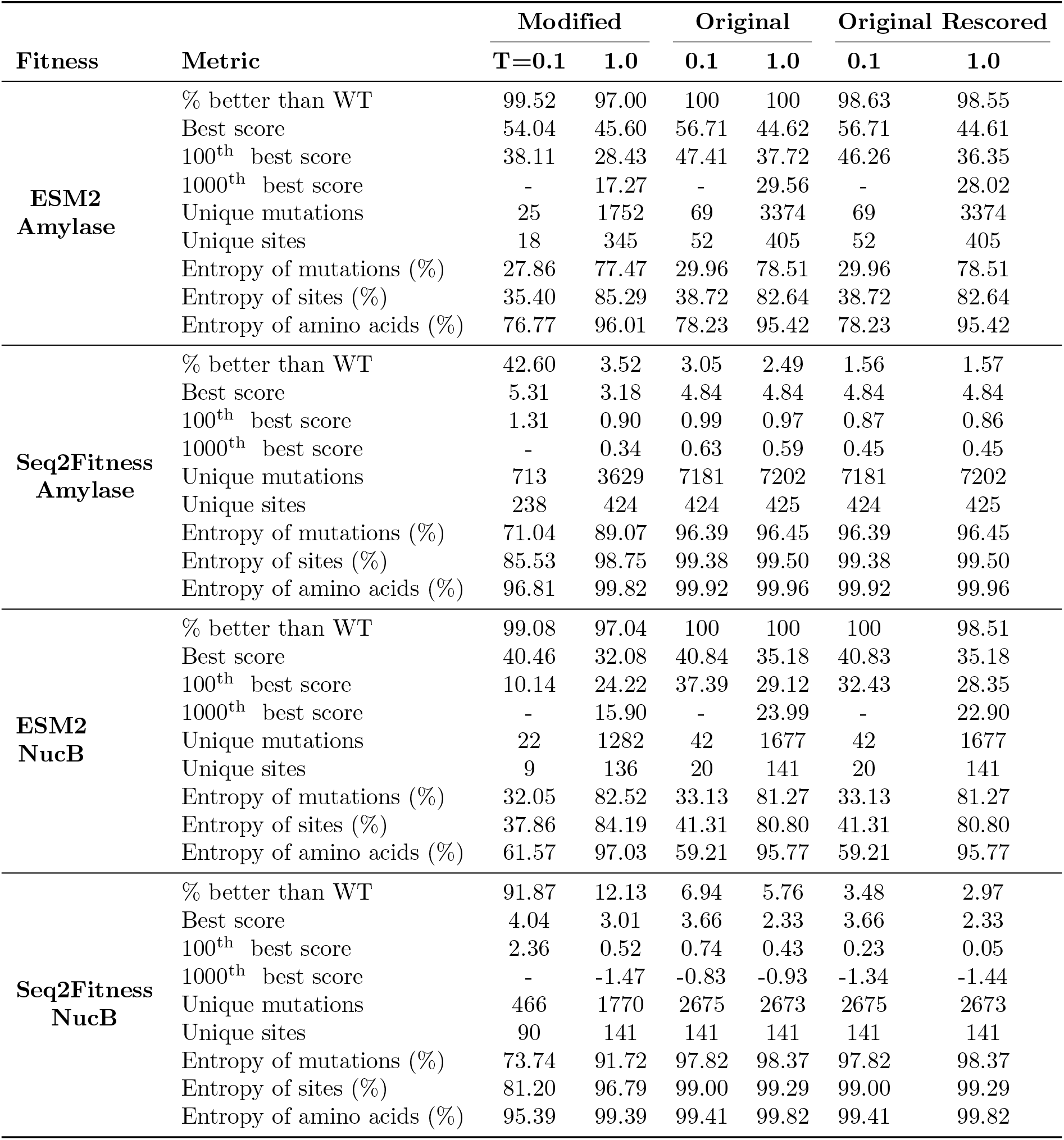
Comparison of EvoProtGrad results obtained using the original EvoProtGrad code and the modified code for designed sequences with only six mutations (k=6). We found that the original code led to inconsistent scores between the optimizer and rescored sequences by the fitness model. We modified the code to ensure consistency.

### The BADASS algorithm

We define the shell of all variants with exactly *k* mutations away from the reference sequence as 𝒮 _*k*_. For a protein with *L* amino acids, this shell contains 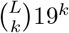 sequences, a number that grows so quickly with *k* that enumerating and scoring all sequences in the shell is only practical for *k* = 1 for typical protein lengths of several hundred amino acids. The BADASS algorithm can explore 𝒮_*k*_, or several shells at once, searching through the larger set ℳ_*k*_ of sequences with *k* or fewer mutations. We let 1 ≤*m* ≤ *M* be an index corresponding to the possible single mutations, where *M* = *L ×*19. We specify an amino acid sequence *x* relative to the reference sequence as the set of mutations in *x*. Let 𝒮_*m*,*t*_ be the set of sequences sampled and scored up to iteration *t* that contain mutation *m*, and 𝒳_*t*_ be the set of sequences sampled at iteration *t*. We describe BADASS next, and discuss how to set its parameters in the Results section.

#### 1. Initialization

Score all *M* single mutant sequences with the fitness model. A reference fitness score *f*_*o*_ and scale *τ* are specified by the user to normalize the scores for numerical stability. We define the normalized energy of mutation *m* as

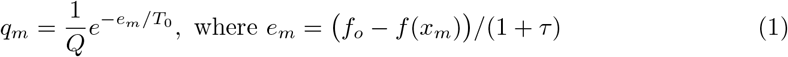

*x*_*m*_ is the sequence with (single) mutation *m, T*_0_ is the initial temperature, and 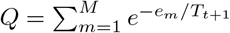 is the partition function. The sequence set is initialized with the single mutants, 𝒮_*m*,0_ = {*f*(*x*_*m*_)}. Then, the high and low average score set point values *µ*_high_ and *µ*_low_ are specified by the user; these values guide the temperature updates. Additionally, the base cooling rate *α*, the heating rate *α*_heat_, and the accelerated cooling rate *α*_cool_ *< α* are defined. The optimizer state is set to *initial transient*.

#### 2. Iteration: Sequences are iteratively sampled until a defined budget is exhausted

a. **Sample sequences**: In each iteration, *N* multi-mutant sequences are sampled from the distribution

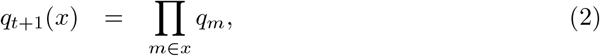

and stored in the set 𝒳_*t*_. Each sequence *x* has *k* mutations when exploring 𝒮_*k*_, or *k* or fewer mutations according to user-specified proportions when exploring ℳ_*k*_.
b. **Score sequences**: Scores for previously sampled sequences in 𝒳_*t*_ are retrieved from a cache, while newly sampled sequences are evaluated using the fitness model. These newly scored sequences are then added to the sets 𝒮_*m*,*t*_, with each new sequence with *k* mutations being added to the *k* sets corresponding to the mutations it contains.
c. **Update optimizer state**: The mean and variance of the fitness scores are computed for sequences sampled during the current iteration, as follows:

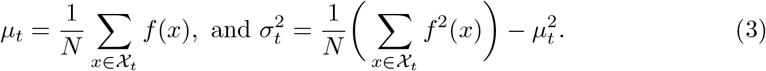

The optimizer state is updated based on the mean score. If the simple moving average of *µ*_*t*_ is greater than *µ*_high_, the optimizer state is set to *active phase transition*. The optimizer remains in this state for a predefined number of iterations (*patience*), or until the score rapidly declines, after which it is switched to *phase transition reversal*. Conversely, if *µ*_*t*_ *< µ*_low_, the state is changed to *cooling phase*.
d. **Update temperature**: The temperature is adjusted based on the current state of the optimizer. If in *initial transient*, the system cools at the base rate, updating the temperature as *T*_*t*+1_ = *αT*_*t*_. Cooling is continued during an active phase transition. However, in *phase transition reversal*, the temperature is increased rapidly according to *T*_*t*+1_ = *α*_heat_*T*_*t*_. During the *cooling phase*, the temperature decreases quickly, following *T*_*t*+1_ = *α*_cool_*T*_*t*_.
e. **Update sampler** through the following calculations:
  i. Mean of mutation scores: 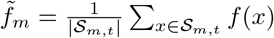
  ii. Variance of mutation scores: 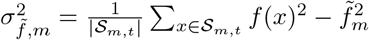
  iii. Raw mutation energies: 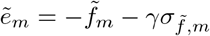 We typically use *γ* = 1.0.
  iv. Normalized mutation energies: 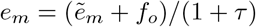
  v. New mutation probabilities: 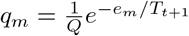

#### 3. Terminate

Once the budget is met, the resulting sequences are analyzed to select the desired ones. The simplest method ranks sequences by their scores and retains the top-ranking sequences. However, more complex selection criteria can be applied, such as limiting the number of sequences with specific mutations or targeting specific mutation sites. In this work, we simplify by retaining only the highest-scoring sequences.

S3 Appendix summarizes BADASS in pseudo-code for easy reference in Figs. 6-8. After an initial transient of tens of iterations, BADASS settles in fairly regular oscillations (e.g., see Fig. 2) that trace out clear patterns in the mean and variance of the model score versus temperature. Figs. 3 and 4 show these traces for the four tasks we consider, along with fits to equations we developed to explain the behavior. For the ESM2 tasks and when sampling sequences with a fixed number of mutations, the change in *µ*_*t*_ and 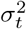 with temperature is monotonic. But for the ML model tasks or the ESM task when we sample a blend of sequences with different numbers of mutations, the variance 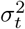 shows an interesting peak at intermediate temperatures. In S4 Appendix, we develop a simple model to understand these behaviors, and state the equations we use to fit the data in Figs. 3 and 4. Next, we motivate BADASS from theory, and study aspects of its convergence to an ideal but impractical sampler.

**Fig 3.**
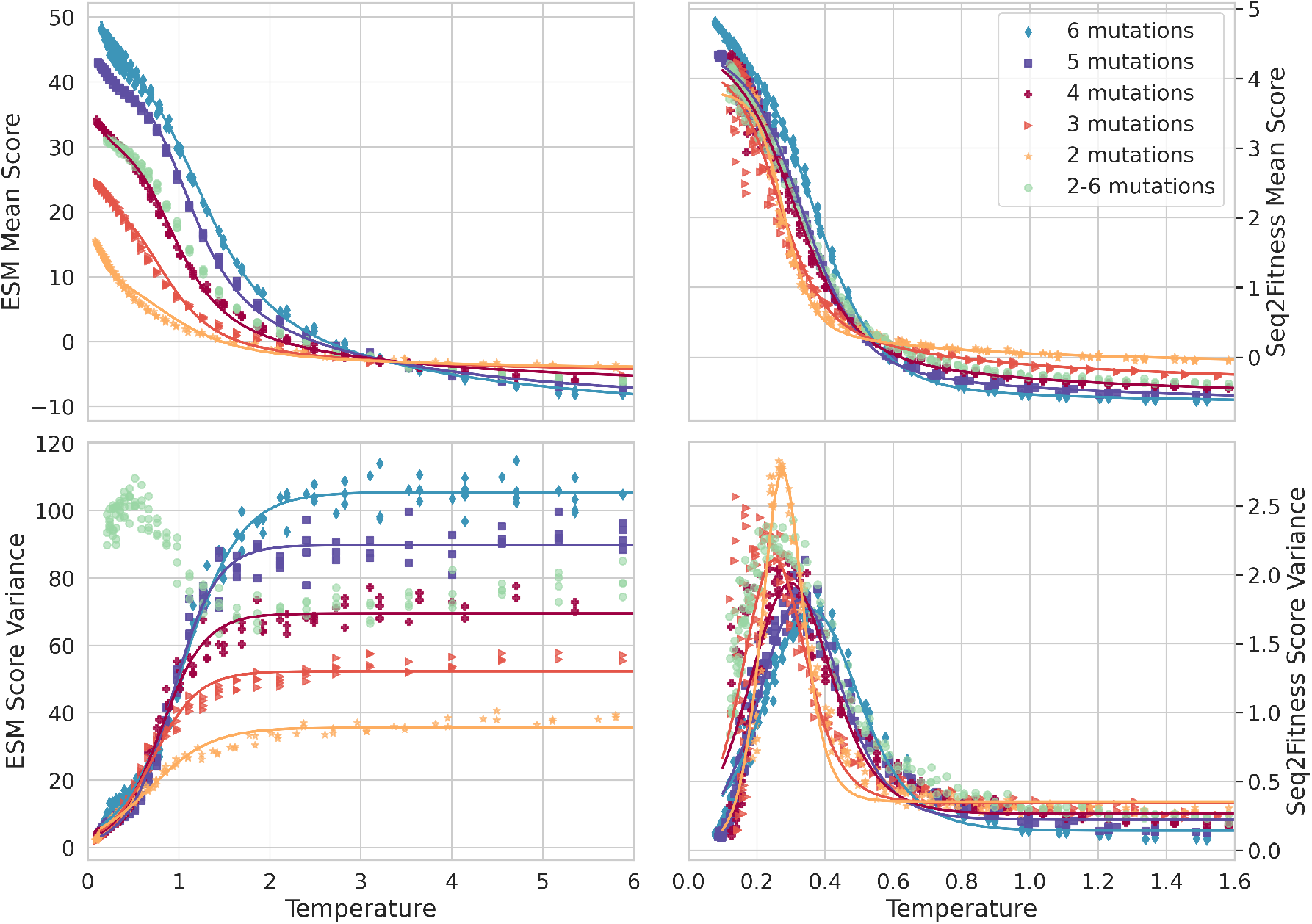
Order parameters versus temperature for amylase tasks: mean score and variance of scores versus temperature after initial transient, i.e., at steady-state oscillatory BADASS behavior. Markers come from BADASS runs, and lines are fits using Eqs. 16, 17 and 18 in S4 Appendix. These were obtained from cooling then heating runs of our algorithm for the amylase task: on the left using the ESM2 mutant marginal score, and on the right using the machine learning model that predicts fitness for stain removal and dp3 function. We ran the algorithm for 250 iterations, scoring 500 sequences in each iteration, and show all data for iterations larger than 100 to avoid the initial transient. The peak of the variance at intermediate temperatures is striking. Running the algorithm with an even blend of numbers of mutations changes the variance behavior, and was not fit to our equations. The mean and variance traces here are reminiscent of the magnetization and susceptibility in Ising models.

**Fig 4.**
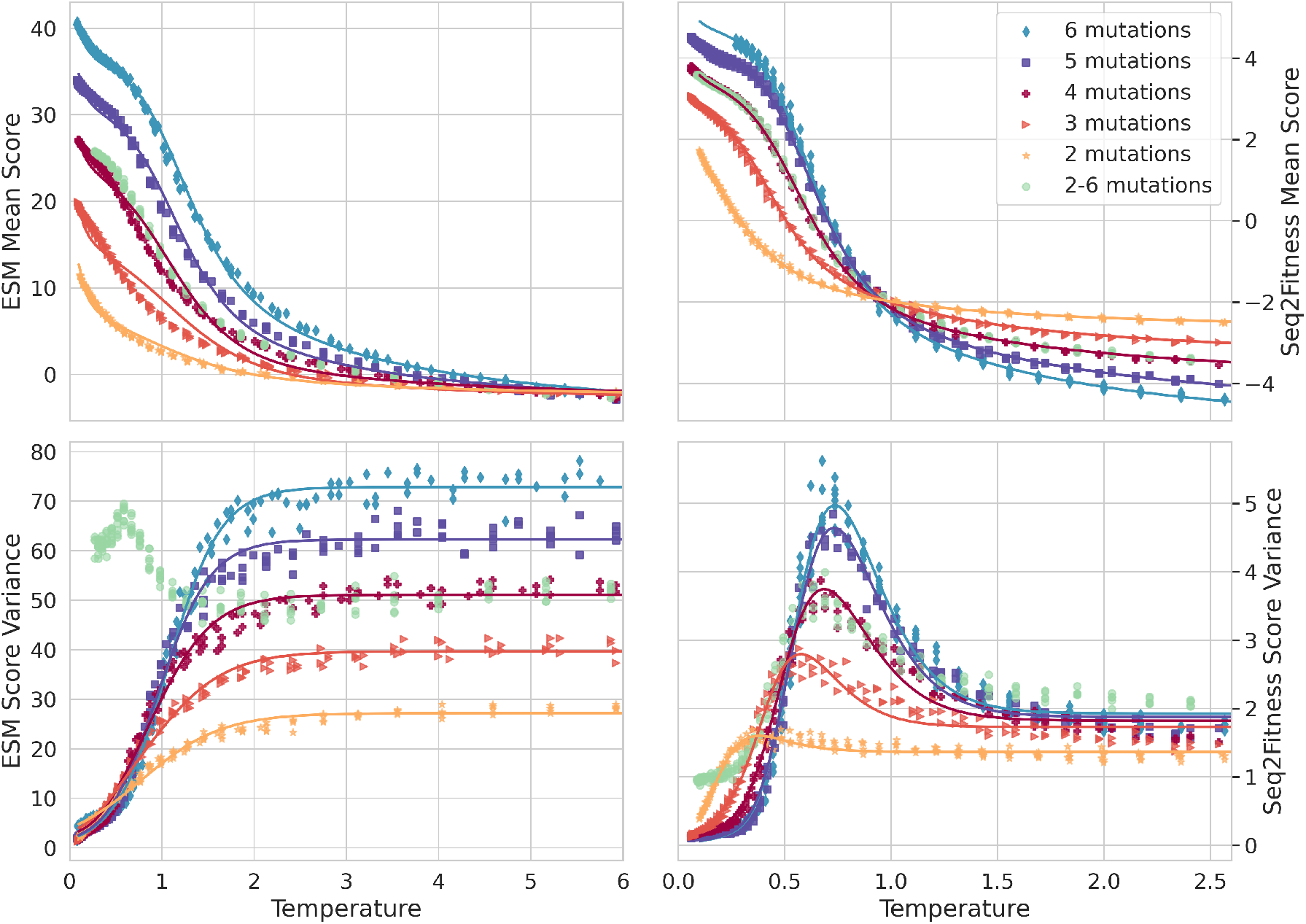
Order parameters versus temperature for the NucB tasks: analogous to Fig. 3. Left corresponds to ESM2 mutant marginal scores, and right the Seq2Fitness model score: the logit for the probability that the nuclease activity is higher than that of the reference sequence.

**Fig 5.**
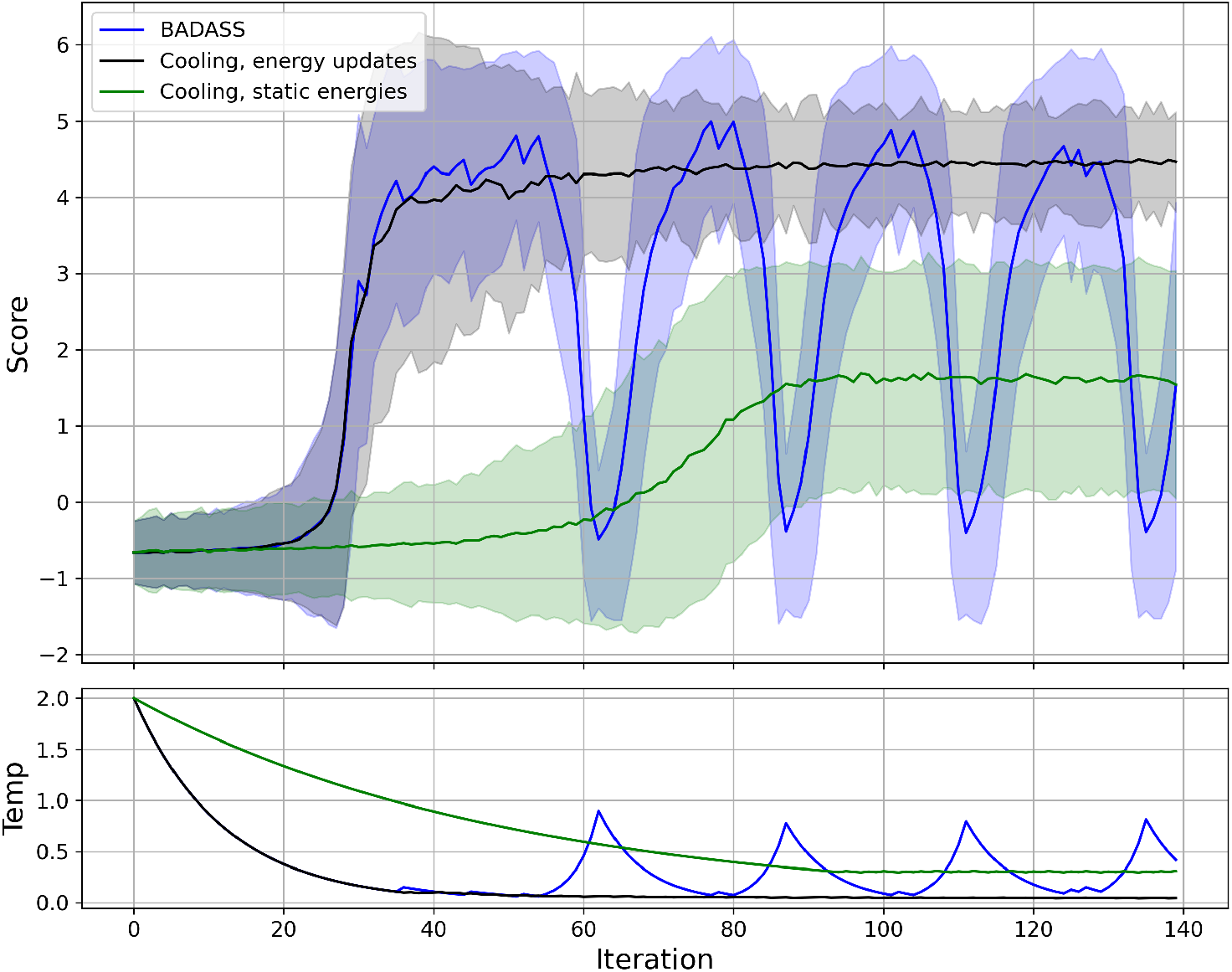
Comparison of BADASS with an approach that simply cools while updating the mutation energies like BADASS does (black), and with an approach that also simply cools and does not update the mutation energies (green) for the Seq2Fitness alpha-amylase task to explore sequences with 6 mutations. Mutation energies for all approaches are initialized to the single mutant scores, with a batch size of 1,000 sequences per iteration. The score envelopes denote the mean *±* 1.96*σ*. The oscillations of BADASS maintain a high variance even when the average score is high, leading to a score envelope that is clearly higher than without the temperature-driven oscillations. Removing the mutation energy updates results in convergence to mediocre scores. The non-BADASS approaches here use minor re-heating when the temperature gets so low that the sampler fails to sample at least half the desired sequence batch size.

**Fig 6.**
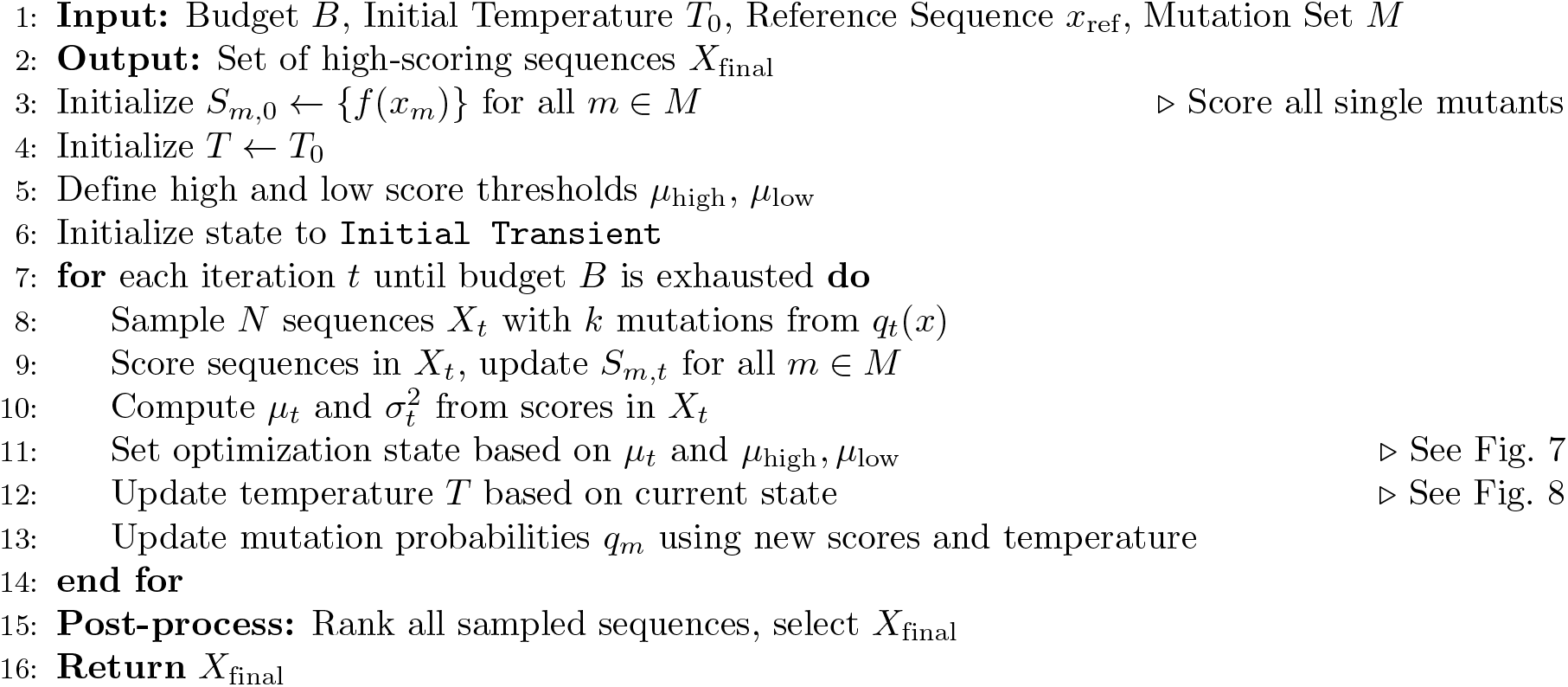
Main Optimization Algorithm.

**Fig 7.**
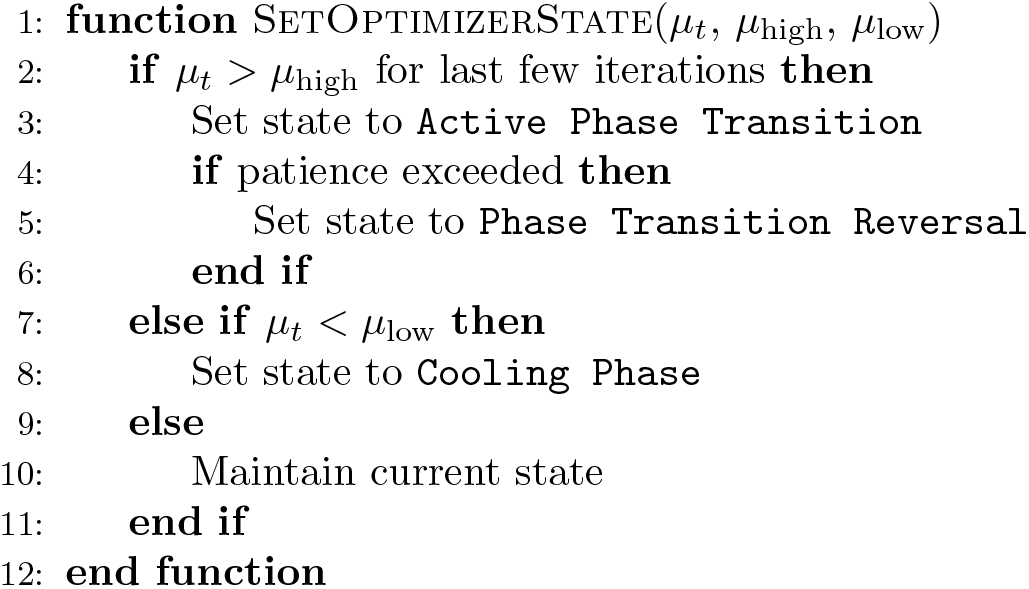
State Logic for the Optimization Algorithm.

**Fig 8.**
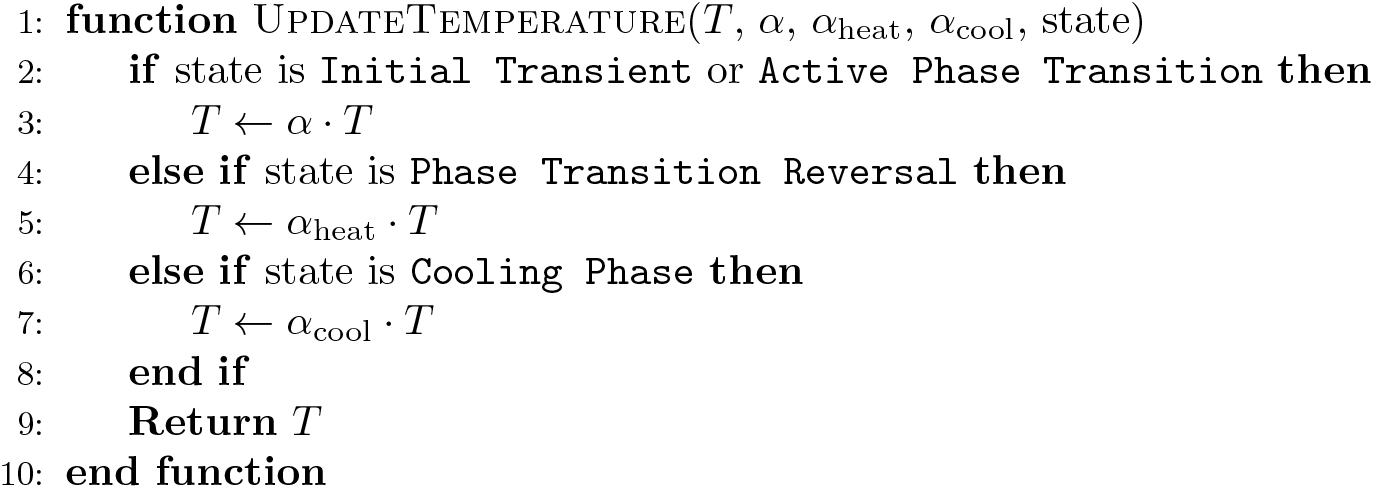
Temperature Update Logic.

### Theoretical motivation for BADASS

To effectively sample diverse, high-scoring sequences, we seek a probability mass function (simply referred to as a distribution going forward) over the sequence space 𝒮_*k*_. Defining 𝒫_*k*_ as the space of such distributions, a natural problem to solve is:

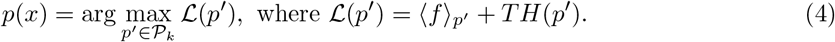

Throughout this work, angled brackets denote averages, with the probability distribution used for the average as a subscript, though omitted when clear from context. So, e.g., ⟨*f*⟩_*p*_*′* =_*x*_ *p*^*′*^(*x*)*f*(*x*). In Eq. 4, *H*(*p*^*′*^) = −⟨log *p*^*′*^(*x*)⟩_*p*_*′* is the Shannon entropy of *p*^*′*^(*x*), and the temperature parameter *T* ≥ 0 controls the trade-off between maximizing the average fitness score and the diversity of *p*^*′*^(*x*). The well-known solution to Eq. 4 is the Boltzmann distribution with energy *E*(*x*) = −*f*(*x*):

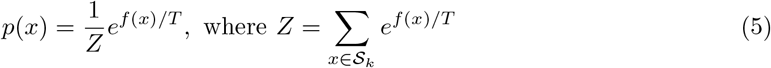

is the partition function. However, because the partition function is the sum of a large number of terms, each requiring evaluation of *f*(*x*), direct computation of *Z* and *p*(*x*) is intractable. Recent approaches enable reasonably effective MCMC sampling from Eq. 5 by defining a tractable proposal distribution with high acceptance probabilities. Specifically, EvoProtGrad [15] introduced a proposal distribution based on the gradients of *f*(*x*) with respect to the amino acids as one-hot encodings, and a subsequent work called GGS [17] used the same proposal distribution, but applied to a modified model trained on an augmented and smoothed dataset to avoid local optima. We compared these approaches to BADASS in our results.

Our approach is different from MCMC. We restrict the space of probability distributions to a smaller subspace 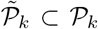 of distributions of the form *q* (*x*) = Π *m* ∈*x* q*m*,where *k* mutations are sampled independently from each other from a distribution over mutations with entries *q*_*m*_ ≥ 0, where1 ≥ *m* ≥ *M* indexes the possible substitutions and 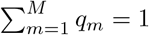 Sampling from any distribution in 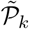 is straightforward, and we can find one that is close enough to Eq. 5 to yield the diverse and high-scoring sequences we seek. To choose the distribution in 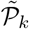, a logical direction is maximizing the same objective as before over the more restricted space, i.e., ℒ (*q*). Rewriting this objective in terms of *q*_*m*_ and calculating its gradient with respect to *q*_*m*_ results in non-linear equations that do not appear to have a closed form solution. But these gradients can be used to optimize ℒ (*q*) numerically with respect to *q*_*m*_, e.g., via Stochastic Gradient Descent as shown recently by [16]. Instead of minimizing ℒ (*p*) directly 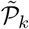,we find the distribution in 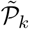 that best approximates the Boltzmann distribution of Eq. 5 in terms of the Kullback-Leibler divergence:

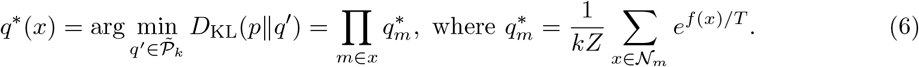

Here we let *𝒩*_*m*_ be the set of all sequences with *k* mutations that contain mutation *m*.^1^

We solve the optimization problem in Eq. 6 to obtain 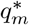 in S5 Appendix. Because 𝒩_*m*_ contains 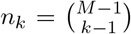 sequences, the entries 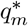 in Eq. 6 cannot be computed in practice, so we make two final approximations and one modification. First, we interpret the summation as proportional to the average of *e*^*f*(*x*)*/T*^ over a uniform distribution over the sequences in *x*∈𝒩_*m*_. We could approximate this average with samples of *x*_*m*_ but the result is statistically inefficient with high variance because of the exponential function, and it leads in practice to poor samplers. So, instead we pursue the mean field approximation

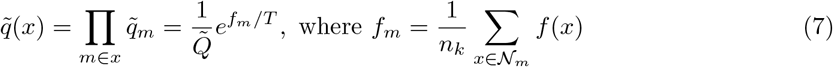

is the average score of all sequences with mutation *m*, and 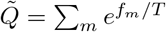 is the partition function. Next, we study how *D*_*KL*_(*p* ∥ *q*^*^) and 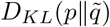 change as the system is cooled by decreasing the temperature. We find that

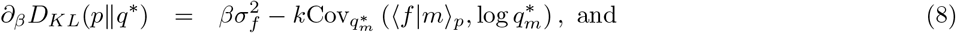

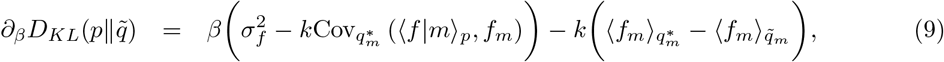

where *β* = 1*/T* is the inverse temperature, ⟨*f* |*m*⟩_*p*_ is the average sequence score when sampling sequences from *p*(*x*) conditioned on the set containing mutation *m*, and 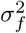 the variance of sequence scores under *p*(*x*). These equations are some of our main results, and are derived in S6 Appendix. For the KL divergences to decrease as the system cools, the covariance terms have to be positive enough for the entire expression to be negative. This can be checked approximately for low and for high temperatures.

We find that at high temperatures, these two KL divergences change approximately at the same small rate proportional to 1*/T*, and have an ambiguous sign. At low-enough temperatures, however, Eq. 8 approximately becomes

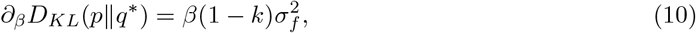

which is negative (assuming a non-zero sequence variance) for *k >* 1, e.g., for sequences with 2 or more mutations, and proportional to 1*/T* . Furthermore, the magnitude increases with *k* as long as 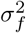 does not decrease faster than linearly with *k* (intuitively, we expect this variance to in fact to grow as *k* increases), predicting a sharper improvement in the closeness between *q*^*^(*x*) and *p*(*x*) as the system is cooled for sequences with an increasing number of mutations, consistent with our empirical results in Figs. 3 and 4, and in the amylase and NucB tables of BADASS results in S2 Appendix. In the case of the mean field approximation 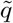, at low enough temperatures,

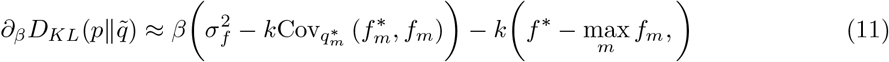

where *f* ^*^ is the maximum score over all sequences, and 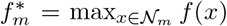 is the largest score of all sequences with mutation *m*. When 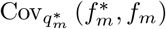 is positive, the KL distance will become smaller as the system cools if the magnitude of the covariance times *k* exceeds 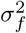, therefore with improved convergence as *k* increases. The low and high temperature approximations of Eqs. 8 and 9 are developed in S7 Appendix and S8 Appendix, respectively. To get to our sampling distribution for BADASS, our second approximation is to estimate *f*_*m*_ using only the sequences we have scored so far that have mutation *m*. This induces a complex bias in our estimate that is optimization-path dependent, but we find that the resulting sampler works well in practice. But the empirical results are often stronger when we encourage the sampling of mutations that have a high standard deviation of scores, in addition to having a large average score, resulting in the modification that yields our final distribution for our sequence sampler defined in step 2e of our algorithm; e.g., compare results in Tables 12 and 14 for *γ* = 0 and *γ* = 1.

## Discussion

We introduced BADASS, a biphasic annealing algorithm that updates mutation energies as it processes sequences, and demonstrated that it consistently outperforms alternative methods for sampling high-scoring protein sequences. While the focus of this work has been on optimizing amino acid sequences, BADASS could be applied to explore other biological sequence spaces, such as DNA, RNA, and even synthetic molecules like lipids and polymers. Additionally, BADASS could be extended to handle more complex mutation types, including insertions and deletions, or multi-output models where multiple properties (e.g., reaction conversion and specificity) must be optimized simultaneously. While MODIFY shares some theoretical similarities with BADASS [16], MODIFY samples mutations from a small subset of carefully selected sites. For design of protein variants where mutations at every site are considered, we observed that MODIFY collapsed and resulted in sequences with substantially lower fitness scores, often failing to identify any sequence with a higher score than the reference. Hence, we focused our comparisons on optimization methods that are designed to handle mutations at all sites.

A major limitation affecting our work, and the broader protein engineering field, is the scarcity of high-quality, large-scale datasets involving multiple mutations, particularly those with detailed functional annotations. Most available datasets focus on single mutations, which limits our ability to train models that generalize to sequences with multiple mutations. Additionally, existing datasets often lack essential metadata, such as experimental conditions like temperature and pH, which is critical for training models that perform consistently across different contexts. To enable further advances in protein design, the field should invest in generating and sharing annotated, multi-mutant datasets across diverse protein families and functions [8]. Collaborative efforts among academic institutions, biotech companies, and public agencies will be crucial in creating these resources. As more standardized multi-mutant datasets become available, they will enable robust benchmarking and the continued development of optimization algorithms like BADASS . To facilitate reproducibility, we have made our code available, encouraging the community to adapt it for their own research.

## Materials and methods

### Seq2Fitness architecture and training

The Seq2Fitness model uses per-residue embeddings from the final transformer layer of the ESM2-650M protein language model to provide rich representations of protein sequences [19]. For each variant, we computed relative embeddings by subtracting the wildtype embedding. We also retrieved logits from the ESM2 output layer and transformed them into log probabilities, resulting in two matrices per sequence: an embedding matrix of size, *L ×* 1280, and a log probability matrix of size, *L×* 20, where L is the sequence length. Each of these matrices were fed to two parallel convolutional paths (Dual-CNN). One path applies a convolution layer followed by a percentile-based statistical summary across the sequence, while the other computes statistical summaries before convolution. Zero-shot fitness scores, including wildtype and mutant marginal scores from ESM2-650M, and masked marginal scores from ESM2-3B [19], as defined by Meier et al [7], were computed as unsupervised fitness predictions. To correct bias in unsupervised scores across sequences with varying number of mutations relative to the wildtype [11], we computed normalized scores by dividing the scores by the number of mutations, and also passed the number of mutations as a feature. The outputs from the convolutional layers and the fitness scores were concatenated and fed to a multi-layer perceptron with two hidden layers. In cases where fitness labels are not comparable, due to variations in screening conditions or assay types, we used multi-task learning with separate outputs for distinct labels and computed the loss as a weighted sum of per-task losses, with identical weights and losses (e.g., sum of squares or cross entropy, depending on the task nature). We standardized fitness labels in the training set to have zero mean and unit variance for numerical stability. We used hyperparameters that included a filter size of 32 for both convolution paths in each Dual-CNN block (Fig. 1B), kernel size of 1, and 11 percentiles for the statistical summaries: 1, 2.5, 12.5, 25, 37.5, 50, 62.5, 75, 87.5, 97.5, and 99. The MLP consisted of two layers with 27 and 15 units, respectively, using the GeLU activation. To mitigate overfitting, we applied a dropout rate of 0.2 and weight decay of 2e-3. The model was trained with an initial learning rate of 1e-2 and a cosine annealing schedule.

We compared Seq2Fitness with various models from the literature. As a baseline, we evaluated zero-shot predictions from ESM2, computed by averaging wildtype and mutant marginal scores [7]. Additionally, we trained an L2-regularized linear model (ridge) with one-hot representations of the proteins (Linear one-hot) and with mean-pooled embeddings from the ESM2 model (Linear ESM) [21]. Following Hsu et al, we also trained augmented ESM models by concatenating one-hot encodings with mutant and wildtype marginal zero-shot scores (Aug. ESM) [10]. For CNN models, we used the architecture proposed by Gelman et al that is most similar to ours (cnn-1xk3f32), which has a kernel size of 3, 32 filters, and a single-hidden layer MLP with 100 units [9, 13]. While their model featurized proteins using one-hot encodings and the top 20 principal components of amino acid index encodings (AAindex) [25], we trained CNNs using this scheme (CNN AAindex), as well as CNNs using only one-hot encoding (CNN one-hot), and per-residue ESM2 embeddings in place of AAindex (CNN ESM).

### Choosing parameters for BADASS

Selecting the right parameters for BADASS involves balancing the robustness of fitness score estimates (which improve with a larger batch size), the frequency of sampling distribution updates (once per iteration, so smaller batch sizes directly imply more sampler updates given a fixed evaluation budget), and GPU utilization when large models like ESM2 or Seq2Fitness are used. Typically, for the protein design tasks discussed here, sampling 500–1000 sequences per iteration strikes a good balance. Our code automatically distributes model inference across available GPUs through PyTorch’s DataParallel, enabling these batch sizes and larger ones if desired for ESM2 and Seq2Fitness models on proteins with hundreds of amino acids. A base cooling rate, *α*, in the range of 0.87–0.94 works well, with heating and accelerated cooling rates, *α*_heat_ and *α*_cool_, set at 1.3–1.8 and *α*^3^ or *α*^2^, respectively. These values help prevent premature convergence while maintaining diversity in the search process. We generally set the reference score *f*_*o*_ to the 80th percentile and set *τ* as the standard deviation of single mutant scores for scaling. Using *γ* = 1 leads to robust results; choosing *γ* = 0 can sometimes produce better sequences, but it can also sometimes do poorly, getting stuck in a subspace with mediocre sequence scores. BADASS typically stabilizes after 60–100 iterations, showing steady oscillations in mean score and variance after that. Running the algorithm for 200–300 iterations often yields a diverse set of high-scoring sequences. With these choices, it takes 15-40 minutes for a single BADASS run on a machine with two NVIDIA RTX4090 GPUs on the design tasks discussed here for budgets between 100,000 and 300,000 sampled sequences. If needed, simulated annealing or an alternative cooling-heating strategy can be accessed via flags in the code, helping to set the score thresholds *µ*_high_ and *µ*_low_ for cooling and heating phases. For different mutation counts, the same general parameter choices tend to work, simplifying tuning across various tasks. The specific parameter values used in this work are available in the code we share.

### Evaluating BADASS

#### Datasets

We conducted protein optimization tasks on two protein families: the alpha-amylase (AMY BACSU) dataset and an internal NucB dataset where fitness is the substrate conversion of a key chemical reaction. The alpha-amylase dataset contained 10,722 unique sequences, each 425 amino acids in length, while the NucB dataset included 55,760 sequences, each with 142 amino acids. These datasets contain multi-mutant sequences, and were chosen to reflect a range of protein sizes and sequence complexity. For both protein families, we evaluated designed sequences containing 2–6 mutations relative to a reference sequence.

#### Tasks

Each optimization task involved finding high-scoring sequences based on two fitness models:

1. **ESM2 Model**: a zero-shot model for scoring protein sequences based on their unsupervised representations. We use the ESM2 650M model, and the mutant marginal score as the fitness. (add equation here?)
2. **Seq2Fitness Model**: a semi-supervised model trained to predict protein function based on experimental data. This model was trained separately for the amylase and NucB tasks. Since fitness labels for NucB are binary, we used the raw logits before sigmoid activation as the optimization metric (i.e. log-probabilities of activity greater than wt labels).

#### Metrics

We compared BADASS to two other sequence optimization methods: EvoProtGrad and GGS. The metrics used to evaluate performance include:

1. **Percentage of sequences better than the wild type**: The proportion of the top 10,000 sequences that improved upon the reference sequence’s score.
2. **Best, best 100th, and best 1**,**000th sequence scores**: Fitness scores of the sequences with ranks 1, 100 and 1,000. Sequences with these ranks or higher would be the prime candidates for wet lab validation.
3. **Unique mutations and mutated sites**: The total number of distinct mutations and sites mutated across the top sequences, reflecting each method’s ability to maintain diversity in the sequence space.

We compare BADASS against two approaches, described next.

**EvoProtGrad** is the original gradient-based Markov Chain Monte Carlo (MCMC)-based method that generates candidate sequences by iteratively proposing mutations and accepting them based on their impact on sequence fitness [15]. EvoProtGrad did better than other competing approaches, but it found far fewer high-fitness sequences than BADASS, particularly in the NucB tasks where it showed poor coverage of sequence space.

**GGS** is a highly intensive, gradient-based MCMC method designed for protein optimization [17] that aims to avoid locally optimal sequence regions by training a new model on an augmented and smoothed dataset. It explores the sequence space by applying small, iterative mutations, following gradients of the fitness function similarly to EvoProtGrad. But GGS also needs to (i) augment and smooth the dataset, (ii) train a sequence to fitness model on the new larger dataset, (iii) use the resulting model to sample sequences with the gradient-based MCMC that is equivalent to EvoProtGrad, and (iv) re-score the sampled sequences with the original model. Because of code compatibility issues, we used EvoProtGrad code in the sequence sampling step for GGS rather than the GGS code. The result is mathematically equivalent to the GGS procedure, with the exception of an additional clustering-based sequence pruning step that we omit. Despite its intensive nature, the results from GGS were not clearly better than BADASS or EvoProtGrad on the original un-smoothed model.

## Supporting information

### S1 Appendix. Seq2Fitness performance on specific datasets and data splits

### S2 Appendix. Detailed results on BADASS amylase and NucB tasks, and comparison to alternate approaches

### S3 Appendix. BADASS algorithm

### S4 Appendix. More on the temperature dependent behavior of the key quantities

Figs. 3 and 4 show how the mean sequence score and variance change versus temperature. We are also interested in a metric called n-effective and denoted by *n*_eff_ that describes how concentrated the sampling distribution is. It is defined as

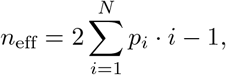

where *p*_*i*_ is a probability mass function with *N* entries sorted from most to least likely. I.e., *n*_eff_ is a scaled average of the rank of the order statistic. We compute three versions: one based on the (joint: sites and amino acid) distribution over mutations, another based on the marginal of the mutation distribution over sites, and the third one based on the analogous marginal over amino acids. Figs. 9 and 10 shows these statistics versus temperature for the same datasets used in Figs. 3 and 4. Changes in *n*_eff_ as BADASS over iterations are often leading indicators of progress, e.g., decreasing as cooling starts many iterations before the average score improves and variance changes.

**Fig 9.**
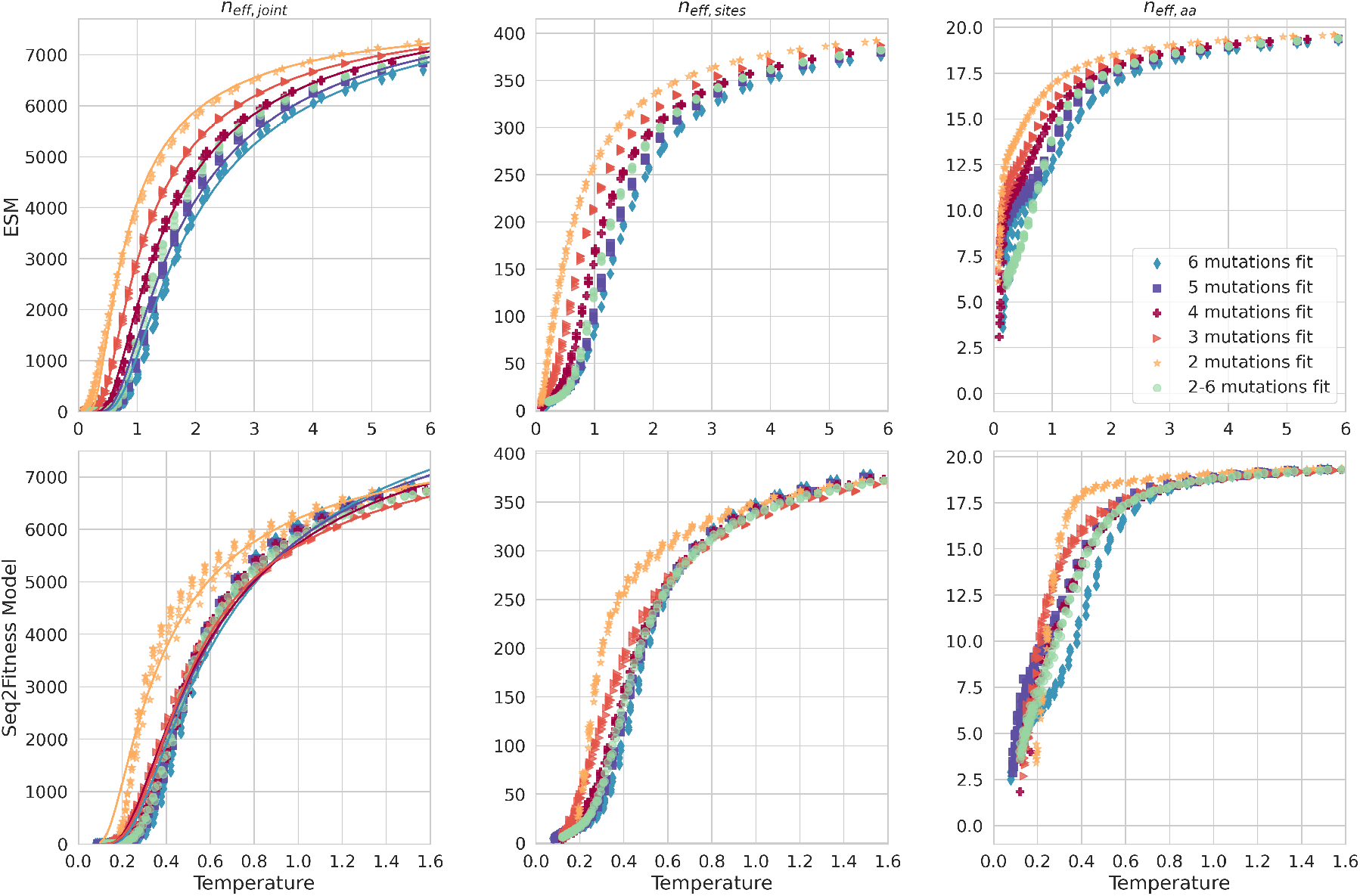
The effective number of levels for the sampling distribution versus temperature for the Alpha Amylase tasks. We also show *n*_eff_ for the marginals over sites and amino acids, and the fit to our equations for the *n*_eff_ version computed from the distribution over mutations (i.e., the joint distribution of sites and amino acids).

**Fig 10.**
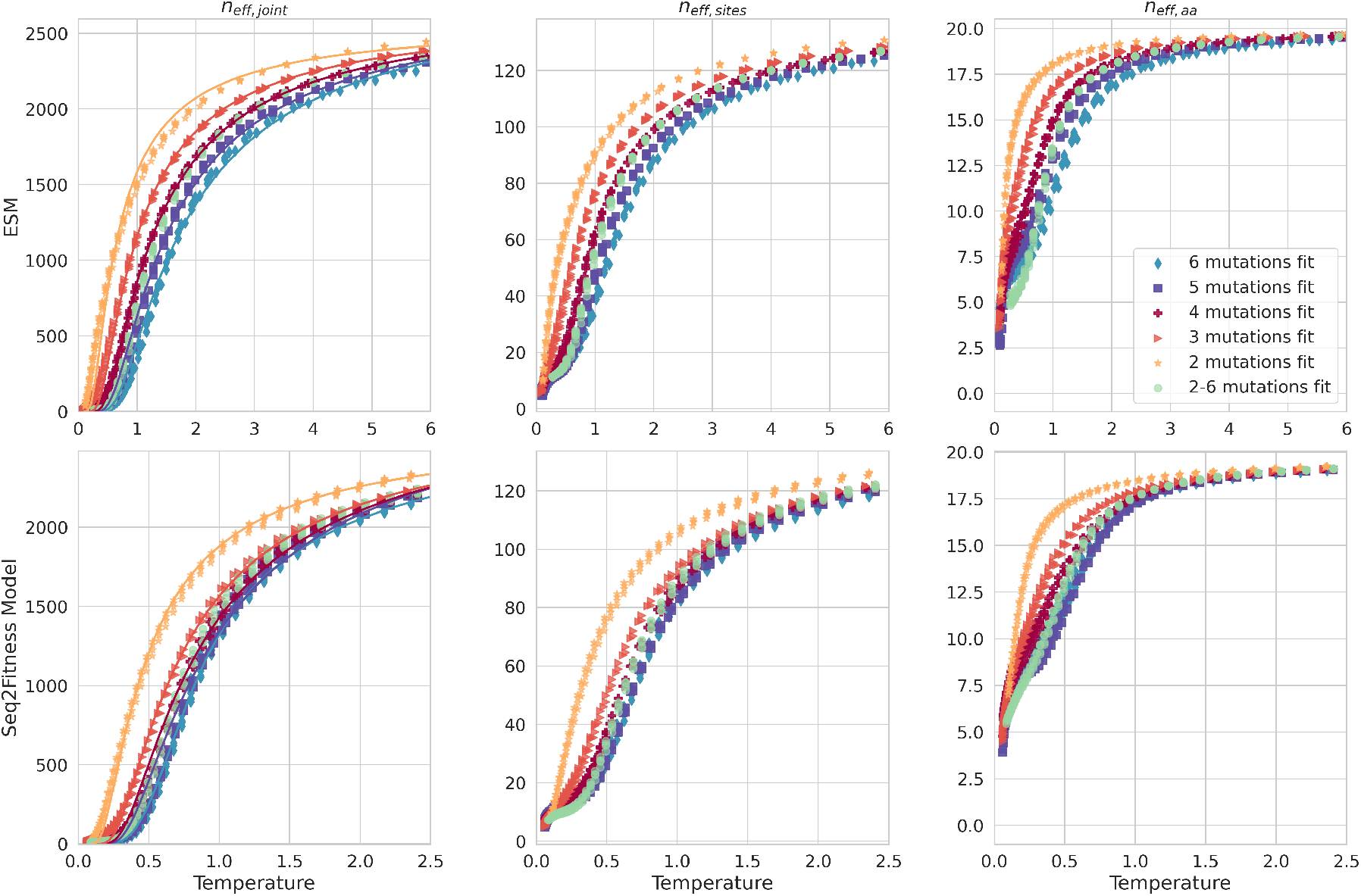
The effective number of levels for the sampling distribution for the NucB tasks.

#### Gaussian Score Model

To understand these behaviors, we consider a simple model where *N* discrete energy levels *E*_*i*_ (negative sequence scores) are sampled i.i.d. from a Gaussian with mean *µ* and variance *σ*^2^. We then define the Boltzmann distribution 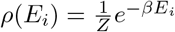 that we use to sample sequences where *β* = 1*/T* is the inverse temperature. So *N* is the size of the sample space we consider, e.g., *N* = *M* for the distribution over mutations. We want to understand the mean, variance and *n*_eff_ for *ρ*(*E*_*i*_). Since the energy levels are Gaussian, we have the following expected value 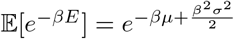.Since *N* is large, we use the mean field approximation

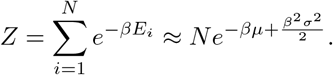

Because of the Boltzmann form of *ρ*(*E*_*i*_), the expected value E[*E*^*m*^*e*^−*βE*^] for any positive integer *m* can be found via:

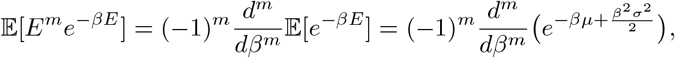

so 𝔼 [*Ee*^−*βE*^] = (*µ − σ*^2^*β*) 𝔼 [*e*^−*βE*^], and 𝔼 [*E*^2^*e*^−*βE*^] = (*µ*^2^ − 2*σ*^2^*µβ* + *σ*^4^*β*^2^ + *σ*^2^) 𝔼 [*e*^−*βE*^]. The average energy becomes

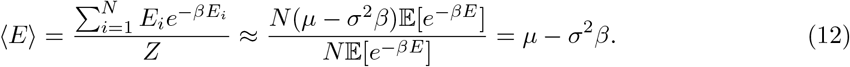

Next, we focus on the variance. A similar mean field approximation of the second moment is ⟨ *E*^2^⟩≈*µ*^2^− 2*σ*^2^*µβ* + *σ*^4^*β*^2^ + ≈*σ*^2^, so Var(*E*) ≈ *σ*^2^. These mean-field results for the average energy and its variance are clearly invalid for large enough *β*, where we know that our distribution will concentrate most of its probability mass in the smallest energy, and have a variance of zero. So we seek a more careful approach when *β* is large next.

For large *β*, the partition function *Z* is dominated by the smallest energies. We use order statistics to approximate the smallest energy levels for a Gaussian distribution. For *N* samples from a Gaussian distribution *N*(*µ, σ*^2^), the i-th order statistic *E*_(*i*)_ can be approximated by:

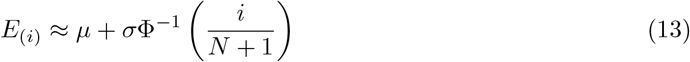

where Φ^−1^ is the inverse cumulative distribution function (CDF) of the standard normal distribution. The smallest two energy levels are:

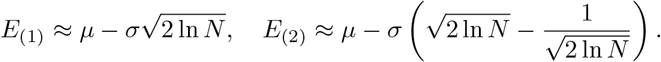

Using the Boltzmann distribution defined on these two energy levels yields after some algebra

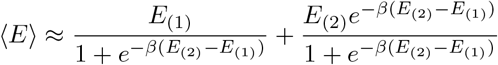

Simplifying since *β* ≫ 1:

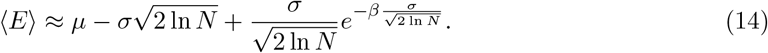

This stays finite as the system freezes, and goes to the expected minimum sample when *N* i.i.d. samples are drawn. Similarly, the second moment becomes

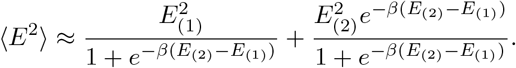

which simplifies when *β* ≫ 1 to:

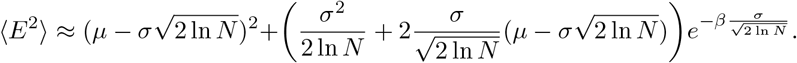

Some algebra and keeping only leading terms in *β* yields

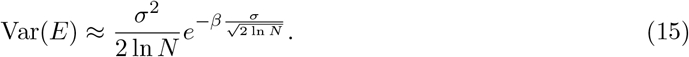

This goes to zero as the system freezes, as expected when only the lowest energy is sampled. To obtain a smooth transition between the mean-field theory and the large *β* behavior using order statistics, we use the interpolating function

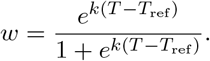

Combining the mean and variance results of the mean field and the large *β* regimes with this choice results in the following functions of *T* :

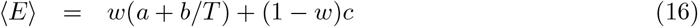

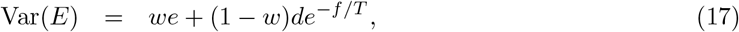

where *a, b, c, d, e, f, k* and *T*_ref_ are scalar parameters. These are the equations we fit to the ESM data in Figs. 3 and 4. For the ML model task, the peak in variance at intermediate temperatures suggests that this simple model is not enough. But a slightly more general one, where the energy labels are sampled from two different Gaussians rather than one, does work. The two modes would correspond to two modes of the sequence scores: one with highly scoring sequences, and the rest.

#### Bi-modal Gaussian Score Model

We now sample the *N* energy levels from the Gaussian mixture with two components 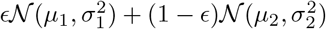.Here, *ϵ* ≪ 1 is the fraction of samples from the first component with a high ML score, and 1 − *ϵ* is the fraction of samples from the second component with a poor ML score. So we assume *µ*_1_ = *µ*_2_− *δ* with *δ >* 0, since we think of energies rather than scores here. Proceeding similarly to the previous model with a single Gaussian component, the mean field approximation for the average energy becomes:

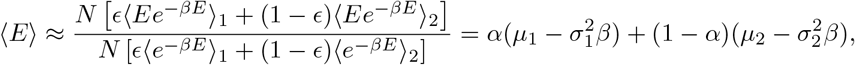

where the subscripts 1 and 2 here denote averages with respect to the two Gaussian components 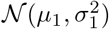 and 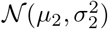, and where

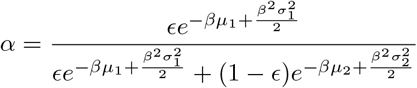

is a sigmoid that transitions between the two modes as a function of temperature. So the average energy is a weighted combination of the average energies of the two modes. To obtain the variance, we first find that the second moment is also the weighted second moment, i.e., ⟨*E*^2^⟩ ≈ *α*⟨*E*^2^⟩_1_ + (1 − *α*)⟨*E*^2^⟩_2_. What is interesting is that subtracting ⟨*E*⟩^2^ to get the variance yields after some algebra:

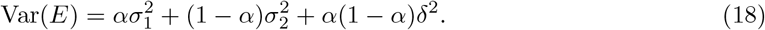

So the variance is not just the weighted variance, and the final additional term explains the variance peak at intermediate temperatures. We also know that at low temperatures, the sigmoid *α* makes the first Gaussian component dominate, so we expect the mean and variance to converge to the same form obtained for the single Gaussian model based on the two lowest energies (of the first Gaussian component). This motivates the single change to the equations we fit to the Seq2Fitness tasks in Figs. 3 and 4 of adding the term *w*(1− *w*)*g* to the variance, where *g* is an additional parameter. More complex functions can be chosen by combining our results here, e.g., using the functional form of *α* for the weight, but the simpler forms suffice to capture the main aspects of the temperature dependent behavior.

#### Temperature dependence of *n*_eff_

Finally, we derive the equation we fit to *n*_eff_ in Figs. 9 and 10. We start with a single Gaussian component model to sample *N* outcomes which are mutations from a Boltzmann distribution with energies *E*_*m*_. So here *E*_*m*_ is the energy that defines the probability of sampling mutation *m*. We have that 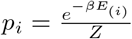 and 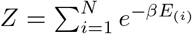, so substitution of Eq.13 yields:

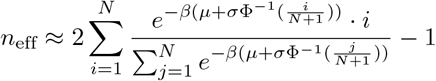

For large *N*, sums can be approximated by integrals. The denominator sum becomes:

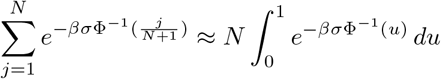

The numerator sum becomes:

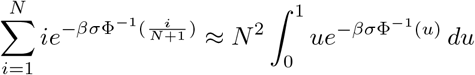

Using the change of variables *v* = Φ^−1^(*u*) and *du* = *ϕ*(*v*) *dv*, where *ϕ*(*v*) is the standard normal density function:

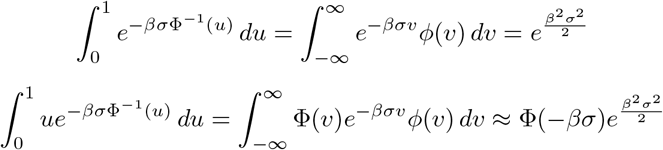

The last integral was approximated by the method of steepest descent. Substituting back into the expression for *n*_eff_:

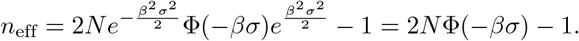

So for small *β*, Φ(−*βσ*) ≈ 0.5 and *n*_eff_ ≈ *N* − 1. For large *β*, using the asymptotic expansion 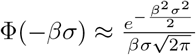 gives 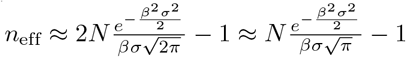. Going through a similar process with the two Gaussian component model for the energies yields:

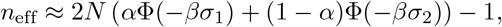

which is just the weighted combination of the per-component *n*_eff_. These results motivate the functions of temperature:

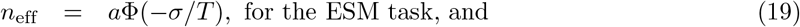

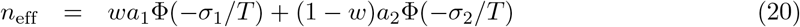

for the Seq2Fitness task. So the first has two parameters, *a* and *σ*, and the second has six (the two sets of *a* and *σ*, and the two parameters *k* and *T*_ref_ for the weight *w*). The resulting fits to the joint distribution of mutations, which is a Boltzmann one, is shown on the left plot of Figs. 9 and 10. These equations also fit the middle plots well, even though the *n*_eff_ is computed over the site marginal distribution, which is not a Boltzmann. The equations do not fit the amino acid version of *n*_eff_ well.

The only remaining piece is the saddle point approximation used above for the integral 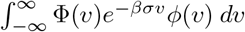 Since 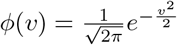, we define the exponent function 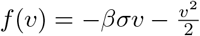 .The saddle point *v*_*0*_ is found by solving 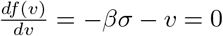,resulting in *v*_0_ = −*βσ*. At the saddle point 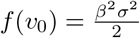 we approximate *f*(*v*) near the saddle point as

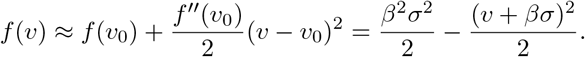

The integral then becomes

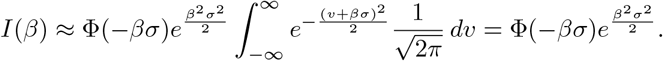

### S5 Appendix. Details Of Sampler Derivation

Here we derive the sampler distribution in Eq. 6. Let 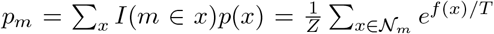 be the probability of sampling from *p*(*x*) a sequence with mutation *m*, where *I*(*m* ∈ *x*) is the indicator function of mutation *m* being in that Σ_*m*_*p*_*m*_ = *k*, so *p*_*m*_ is not a distribution over mutations; *p*_*m*_*/k* is. Substituting the form of *q*(*x*) sequence *x*, and where *N*_*m*_ is the set of sequences with *k* mutations that have mutation *m*. Note into *D*_KL_(*p*∥*q*) and some algebra yields:

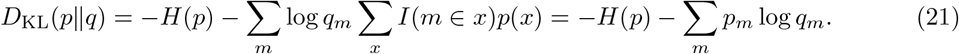

Taking the derivative w.r.t. *q*_*m*_ and setting it to zero gives

The first term is the negative entropy of *p*(*x*), which is independent of *q*_*m*_ and can be ignored in the optimization. The second term is the cross entropy. To minimize it, we define the objective where we add a Lagrangian multiplier to ensure the result adds up to 1 over mutations. 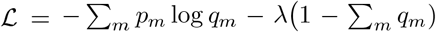. Taking the derivative w.r.t. *qm* and setting it to zero gives 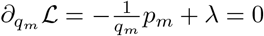. Solving for *q*_*m*_, normalizing its sum over mutations to one, and noting that 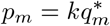 completes the derivation of Eq. 6. Substituting 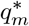 back in the cross entropy term, we find it is equal to 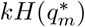,i.e., *k* times the entropy of 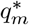. So 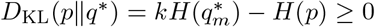. Also, any other 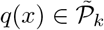 will have a larger (or equal) *D*_KL_(*p*∥*q*) than 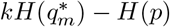.

### S6 Appendix. Convergence of Sampler as Temperature Drops

Here we study how *D*_KL_(*p*∥*q*) changes as the system is cooled for the two key distributions in the derivation of our sampler: *q*^*^(*x*), and its mean field approximation 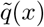.These sampling distributions use the scores of all possible sequences, an assumption that makes our analysis here tractable but limits its applicability to our actual sampling distribution which approximates 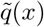 with available samples. A convergence analysis for our actual sampler remains an open problem. Our goal is understanding how *D*_KL_(*p*∥*q*) changes as the inverse temperature *β* = 1*/T* increases. For any distribution 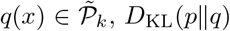,breaks up into the two terms in Eq. 21. We derive the derivative with respect to *β* for the two terms separately. Starting with the entropy term, we substitute *p*(*x*) from Eq. 5 to find

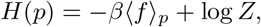

where ⟨*g*⟩_*p*_ denotes the average of any function *g*(*x*) over the distribution *p*(*x*), a compact notation we will use throughout this section. Simple algebra shows that the expectation of the energy is ⟨*f*⟩_*p*_ = ∂_*β*_ log *Z*, and that the second derivative of log *Z* with respect to *β* is 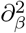 log 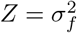,where 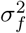 is the variance of sequence scores under *p*(*x*). The latter implies 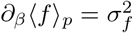.So taking the derivative of the entropy *H*(*p*) with respect to *β* yields

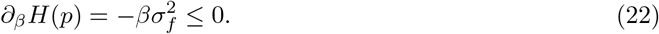

I.e., the entropy of *p*(*x*) can only decrease as the system cools, matching intuition. Next we focus on the second term in Eq. 21, which we denote here by 𝓁 =_*m*_*p*_*m*_ log *q*_*m*_. Substituting 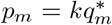 into 𝓁, we obtain

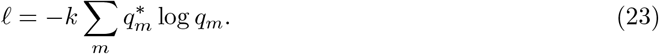

So the cross entropy term in *D*_KL_(*p*∥*q*), where the arguments are distributions over sequences, simplifies into *k* times the cross entropy between the optimal mutation distribution 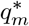 and the so-far arbitrary mutation distribution *q*_*m*_. Next, some algebra results in

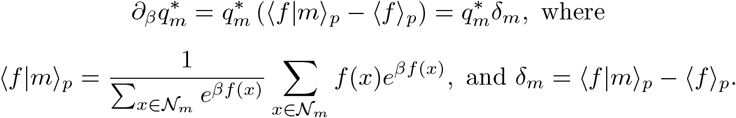

I.e., ⟨f|m⟩*p* is the conditional average sequence score under p(x) when restricting the sequence set to those with mutation *m*. Note that 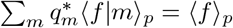 so 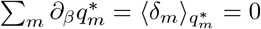 . Moving on, we take the derivative of 𝓁:

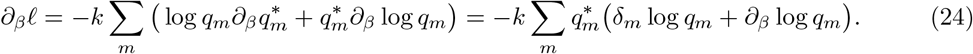

We are ready to determine this derivative for our two sampling distributions. We start with 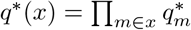. Since the above implies that 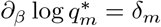,we substitute 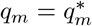 in Eq. 24 to find that

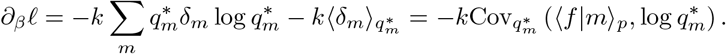

Combining this derivative with the one for the entropy in Eq. 22 yields the expression for the change in *D*_KL_(*p*∥*q*^*^) with respect to *β* in Eq. 8. Next, we work on the derivative of 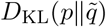.When 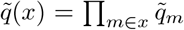 we have that log 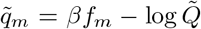, and 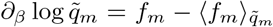.Substituting these expressions in Eq. 24 and some algebra yields

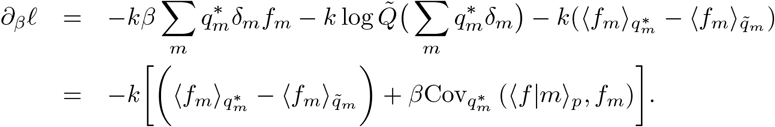

Combining this derivative with the entropy term derivative in Eq. 22 yields the derivative of 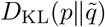 with respect to *β* in Eq. 9.

### S7 Appendix. Low temperature approximation of convergence results

Our goal is to approximate Eqs.8 and 9 when *β* ≫ 1 to leading order. Our main approximation is that sums of Boltzmann factors become concentrated on the largest one exponentially fast. Assuming there is a single sequence that attains the maximum score in 𝒩_*m*_ for all *m* to simplify, and letting 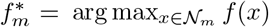 be the largest sequence score from the set *N*_*m*_, we approximate 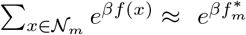 to get

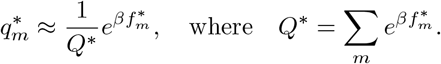

So 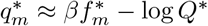. The approximation 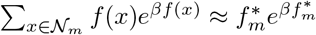 yields 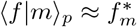 So

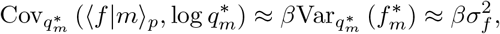

The last approximation follows from the law of total variance

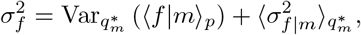

where the last term is zero to leading order because within 𝒩_*m*_ the probability of sequences gets concentrated in 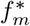.Substituting this approximation in Eq. 8 we obtain Eq. 10. Next we work on the approximation for Eq. 9. Using the approximation for ⟨*f*|*m*⟩_*p*_, we find that

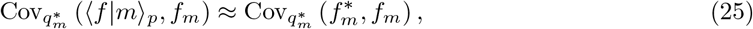

which is bounded in magnitude above through Cauchy-Schwartz by

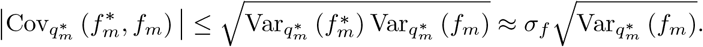

But we cannot express this more fully in terms of 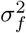.Now we tackle the second term in Eq. 9, which involves the difference of averages of *f*_*m*_. We approximate 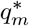 further now to only include the top two values of 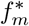, which we denote by 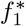,and 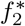So

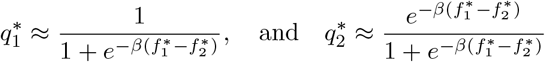

for the two mutations with the two highest 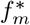,while other entries of 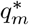 are zero. We similarly approximate 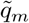 with the two largest values of *f*_*m*_:

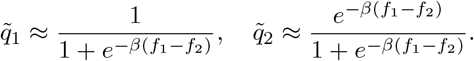

Using these approximations, we compute the averages

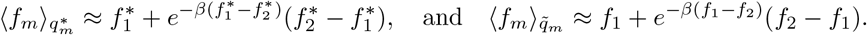

Taking their difference

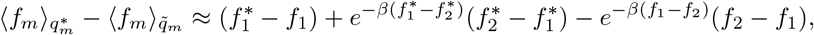

which is non-negative for large enough *β* since 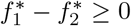 . In the large *β* limit, the difference is dominated by the term 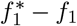, with exponentially small corrections due to the second-largest values 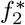 and *f*_2_. This shows the second term in Eq. 9 is smaller than zero in this limit. Substituting our results for both terms in the equation, we obtain Eq. 11.

### S8 Appendix. High temperature approximation of convergence results

Here we analyze the behavior of Eqs. 8 and 9 at high temperatures, with *β* ≪ 1. We approximate the necessary quantities to first order in *β*, mostly relying on the first-order Taylor approximation *e*^*βf*(*x*)^ ≈ 1+*βf*(*x*). Applying this to 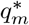, its numerator becomes 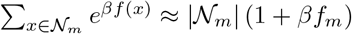 Similarly, the partition function is 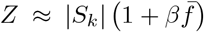, where 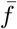 is the uniform average score over all sequences in *S*_*k*_. Substituting these into the expression for 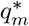, and noting that 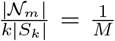, where *M* is the total number of mutations, we get

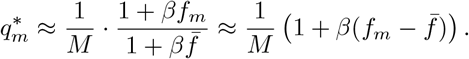

Taking the logarithm of 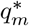 and using the approximation log(1 + *x*) ≈ *x* for small *x* to obtain 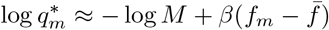. Next, we focus on the conditional expectation ⟨*f*|*m*⟩_*p*_. Substituting the first-order Taylor expansion of *e*^*βf*(*x*)^ in the numerator and denominator of its definition, we have

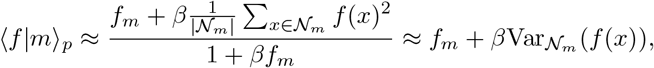

where the variance is based on uniform sampling of sequences in *𝒩*_*m*_. Substituting the approximations for log 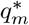 and ⟨*f*|*m*⟩_*p*_ into the covariance term in Eq. 8, we have

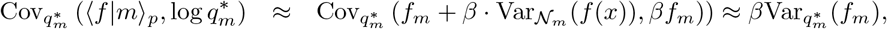

to leading order in *β*. Using our approximation above for 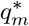, we approximate this variance next. First, we approximate the expectations:

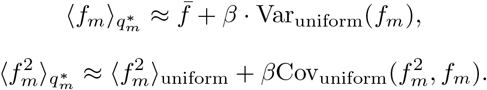

Substituting these into the variance expression and expanding to first order in *β*, we get

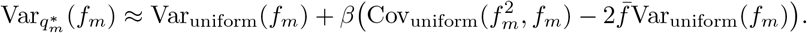

Using the analogous small-*β* approximation for the Boltzmann distribution 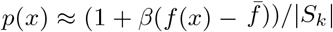, the variance of scores 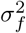 under *p*(*x*) can be expressed as:

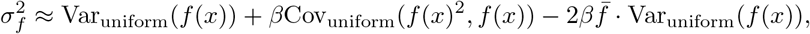

where Var_uniform_(*f*(*x*)) is the variance of *f*(*x*) under uniform sampling of sequences, and 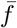 is the uniform average of *f*(*x*) over all sequences. So to leading order, Eq. 8 becomes

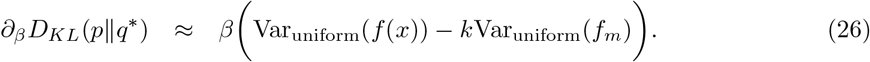

This is small in magnitude because of *β* and has an ambigious sign: if Var_uniform_(*f*_*m*_) is small then the expression is positive, e.g., if Var_uniform_(*f*_*m*_) ≈ Var_uniform_(*f*(*x*))*/*| *𝒩*_*m*_|, if the scores for all sequences are approximately random. If Var_uniform_(*f*_*m*_) is large, e.g., if the sequences of scores in the same set *𝒩*_*m*_ are positively correlated, the expression can be negative. Next, we use the same approximations to analyze ∂_*β*_*D*_*KL*_(*p*∥*q*^*^) as described in Eq. 9. Substituting the approximations, we find that

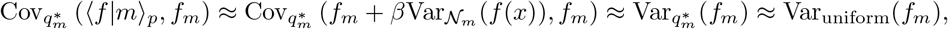

where we drop all terms linear or higher in *β*, since that is all we need to recover terms to first order in Eq. 9. The small *β* approximation of 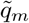 ends up being identical to that of 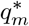 up to first order in *β*, so the second term in Eq. 9 is zero at this level of accuracy. Substituting the approximation above for 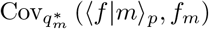 in the first term yields 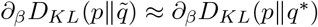.

## Acknowledgments

We are grateful to Solugen for supporting this work. Yuriy Roman-Leshkov carefully read a draft of this work, and provided valuable suggestions. We thank Patrick Emami for insightful discussions about the EvoProtGrad approach and code.

## Footnotes

1 Substituting *q*^*^(*x*) into *D*_KL_(*p*∥*q*) yields the non-trivial inequality 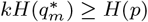 that provides an upper bound of the entropy of the Boltzmann distribution in Eq. 5 in terms of the the entropy of the optimal distribution over mutations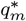 . It is valid for any positive integer *k*, although *p*(*x*) and 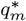 also depend on *k*; the former through its support, and the latter through the sets 𝒩_*m*_.

